# Bat pluripotent stem cells reveal unique entanglement between host and viruses

**DOI:** 10.1101/2022.09.23.509261

**Authors:** Marion Déjosez, Arturo Marin, Graham M. Hughes, Ariadna E. Morales, Carlos Godoy-Parejo, Jonathan Gray, Yiren Qin, Arun A. Singh, Hui Xu, Javier Juste, Carlos Ibáñez, Kris M. White, Romel Rosales, Nancy J. Francoeur, Robert P. Sebra, Dominic Alcock, Sébastien J. Puechmaille, Andrzej Pastusiak, Simon D.W. Frost, Michael Hiller, Richard A. Young, Emma C. Teeling, Adolfo García-Sastre, Thomas P. Zwaka

## Abstract

Bats have evolved features unique amongst mammals, including flight, laryngeal echolocation, and certain species have been shown to have a unique immune response that may enable them to tolerate viruses such as SARS-CoVs, MERS-CoVs, Nipah, and Marburg viruses. Robust cellular models have yet to be developed for bats, hindering our ability to further understand their special biology and handling of viral pathogens. To establish bats as new model study species, we generated induced pluripotent stem cells (iPSCs) from a wild greater horseshoe bat (*Rhinolophus ferrumequinum*) using a modified Yamanaka protocol. Rhinolophids are amongst the longest living bat species and are asymptomatic carriers of coronaviruses, including one of the viruses most closely related to SARS-CoV-2. Bat induced pluripotent stem (BiPS) cells were stable in culture, readily differentiated into all three germ layers, and formed complex embryoid bodies, including organoids. The BiPS cells were found to have a core pluripotency gene expression program similar to that of other species, but it also resembled that of cells attacked by viruses. The BiPS cells produced a rich set of diverse endogenized viral sequences and in particular retroviruses. We further validated our protocol by developing iPS cells from an evolutionary distant bat species *Myotis myotis* (greater mouse-eared bat) non-lethally sampled in the wild, which exhibited similar attributes to the greater horseshoe bat iPS cells, suggesting that this unique pluripotent state evolved in the ancestral bat lineage. Although previous studies have suggested that bats have developed powerful strategies to tame their inflammatory response, our results argue that they have also evolved mechanisms to accommodate a substantial load of endogenous viral sequences and suggest that the natural history of bats and viruses is more profoundly intertwined than previously thought. Further study of bat iPS cells and their differentiated progeny should advance our understanding of the role bats play as virus hosts, provide a novel method of disease surveillance, and enable the functional studies required to ascertain the molecular basis of bats’ unique traits.

## Introduction

It has been an age-old question: what makes bats so fascinating to humans? Bats account for one-fifth of all living mammalian species (*n*=1,451) (Simmons and Cirranello, 2020), inhabiting diverse ecological niches, feeding on arthropods, fruit, nectar, leaves, fish, blood, and small vertebrates (Allen, 1939; Nagel, 1974; Teeling et al., 2012; Teeling et al., 2018). They are the only mammals to have evolved true, self-powered flight and can use laryngeal echolocation to orient in complete darkness (Teeling *et al*., 2012; Teeling et al., 2005). They are found throughout the globe, absent only from extreme polar regions (Altringham, 2011; Teeling *et al*., 2012; Teeling *et al*., 2005). Many bat species studied exhibit an extremely long lifespan relative to body size and a suspected low tumorigenesis rate (Munshi-South and Wilkinson, 2010; Wilkinson and Adams, 2019) (see ref (Wilkinson and Adams, 2019) for review). Still, what makes bats most distinctive is that many species (e.g., rhinolophids, hipposiderids, pteropodids) have been shown to contain some of the richest virospheres amongst mammals (Hermida Lorenzo et al., 2021; Luis et al., 2013; Olival et al., 2017) including the closest known relatives of SARS-CoV, SARS-CoV-2, MERS-CoV, Marburg and henipaviruses (Anthony et al., 2017; Brook and Dobson, 2015; Drexler et al., 2012; O’shea et al., 2014; Olival *et al*., 2017; Van Brussel and Holmes, 2022; Wang et al., 2011). This is potentially due to a modulation of their innate immune response rendering them as asymptomatic and tolerant viral hosts (Banerjee et al., 2020; Brook et al., 2020; Pourrut et al., 2009; Yob et al., 2001). Also, despite being among the smallest mammalian genomes, bat genomes contain the highest diversity of ancient and contemporary viral insertions of retroviral and non-retroviral origin (Cui et al., 2012; Hayward et al., 2018; Jebb et al., 2020; Skirmuntt et al., 2020; Skirmuntt and Katzourakis, 2019), suggesting that bats have a long and tolerant evolutionary history with their viral pathogens. As some of the integrated retroviral sequences are full length and even of non-bat origin, sequencing bat genomes provides novel insights into the bat virosphere and the potential for zoonotic spillover (Cui et al., 2015; Drexler *et al*., 2012; Escalera-Zamudio et al., 2016; Hayman et al., 2013), but also uncover mechanisms of viral persistence (Skirmuntt *et al*., 2020). To date, how bats deal with viruses is still poorly understood with only a limited number of immune cells documented and characterized in a few bat species (e.g., *Pteropus alecto, Eonycteris spelaea, Myotis lucifugus, Eptesicus fuscus*) (Middleton et al., 2007; Swanepoel et al., 1996; Watanabe et al., 2010). Further, developing novel cellular resources and assays is required to uncover and validate the molecular adaptations that have evolved in bats to tolerate these viral pathogens (Jebb *et al*., 2020; Middleton *et al*., 2007; Swanepoel *et al*., 1996; Watanabe *et al*., 2010). The prevailing hypothesis supported by recent comparative genomic analyses of multiple bat families (Santillán et al., 2021) is that viral tolerance results from specific adaptations of their innate immune system. Accordingly, bats mount an inaugural antiviral reaction after viral inoculation like all mammals but then quickly “dampen” this very response before it becomes overly pathological (Jebb *et al*., 2020; Kacprzyk et al., 2017; Wang et al., 2021). Critically, this unusual way to deal with viruses could be caused in part by molecular adaptations that stifle canonical virus sensing and the subsequent inflammatory response (Baker et al., 2013; Banerjee *et al*., 2020; Irving et al., 2021; Wang *et al*., 2011) such as the cGAS-STING, OAS-RNASE L, and NLPR3 systems (Ahn et al., 2019; Goh et al., 2020; Mozzi et al., 2015; Pavlovich et al., 2018; Xie et al., 2018).

It is striking how closely the aforementioned genomic adaptations to the bat immune system mirror how viruses themselves typically dismantle the host response (García-Sastre, 2017). Yet viruses do not just use countermeasures against detection and quench inflammation. Viruses are also infinite masters of tweaking cell processes to their advantage to convert host cells into virus-producing factories. Hence, we wondered if, in addition to the immune evasion strategies of viruses, bats also harbor the blueprints for productive viral replication, given the evolutionary maintenance of many intact and full-length viral elements in one of the smallest mammalian genomes, and the potential evolutionary advantage gained from such a symbiosis (Villarreal et al., 2000; Witzany, 2010).

Here, we sought to test empirically the idea that bats genetically simulate the viral ploy for immune evasion and promote notably fertile ground for virus production. We conjectured that pluripotent stem cells would be an ideal experimental system for addressing this question. Given that pluripotent stem cells are the founding cells of the entire embryo, their cellular ground state provides an exclusive reference point for comparative studies as all mammals must complete this stage in a similar manner (Wray et al., 2010). Importantly, the global epigenetic resetting that occurs as cells reprogram to pluripotency causes the transcriptional reactivation of endogenous viruses (Grow et al., 2015; Macfarlan et al., 2012; Rowe and Trono, 2011; Wang et al., 2014). As such, it would present a unique window into the abundant endogenized viral diversity within bat genomes, allowing the broad cataloging of active viruses and, in turn, the study of how viruses interface with host cell programs.

## Results

### Bat Yamanaka Reprogramming

Given the importance of bats as an emerging model system in multiple areas and the need to study their unique biology and potential role in pandemics, we sought to develop an effective strategy to produce bat pluripotent stem cells (Figure 1A). Despite numerous attempts by different groups (Mo et al., 2014; Wang *et al*., 2021), and some initial success in partial reprograming (Aurine et al., 2021), robust pluripotent stem cell lines from bat species, with globally repeatable protocols, have not been previously reported.

**Figure 1.**
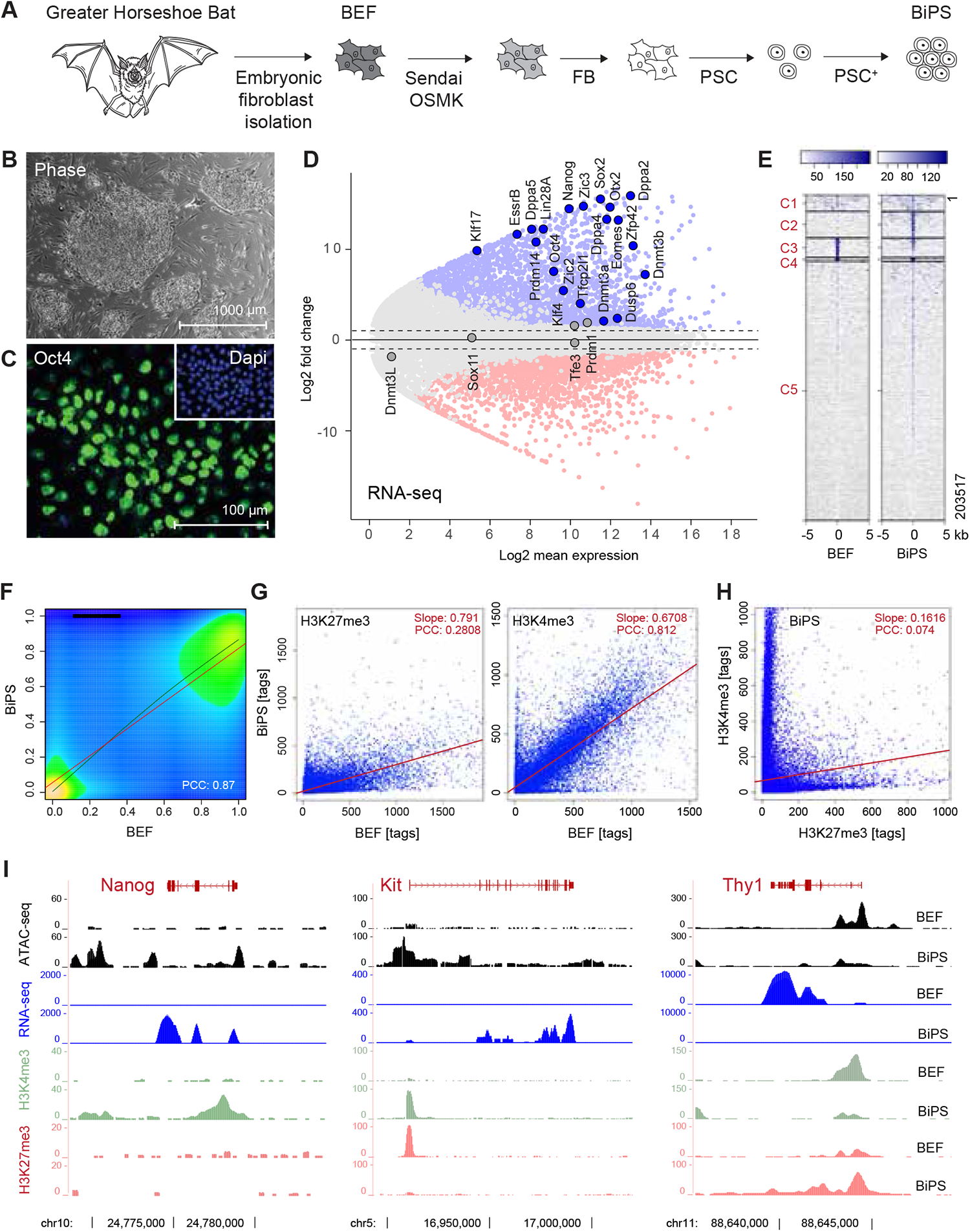
Derivation of pluripotent bat stem cells. **(A)** Illustration of the bat pluripotent stem cell derivation strategy. BEF, embryonic fibroblasts; OSMK, Oct4, Sox2, cMyc, Klf4; FB, fibroblast medium; PSC, pluripotent stem cell medium; PSC+, PSC with additives. **(B)** Morphology of established BiPS cell colonies grown on mouse embryonic fibroblasts. **(C)** Immunofluorescent detection of Oct4 in BiPS cells. **(D)** MA plot of RNA-seq data illustrating the transcriptional differences between bat embryonic fibroblast (BEF) and pluripotent stem cells (BiPS). Selected genes with known functions in the establishment or maintenance of pluripotency are highlighted. **(E)** Kmean cluster analysis of ATAC-seq signals obtained from BEF or BiPS cells. C, cluster. **(F),** Density plot of RRBS results obtained from BEF and BiPS cells. PCC, Pearson correlation coefficient. **(G)** Scatter plots of histone 3 methylation status at K4 (activating chromatin modification) or K27 (repressing chromatin modification) after ChIP-seq from BEF or BiPS cells as indicated. **(H)** Scatter plot of H3K4me3 and H3K27me3 in BiPS cells illustrating the occurrence of bivalent chromatin sites in BiPS cells. **(I)** RNA-seq, ATAC-seq and H3K4me3 or H3K27me3 ChIP-seq signals of selected genes with known roles in reprogramming that are activated (Nanog, Kit) or repressed (Thy1) in BiPS when compared to BEF cells.

We first focused on the original Yamanaka reprogramming paradigm based on four reprogramming factors (Oct4, Sox2, Klf4, and cMyc) because it provides the most direct way to generate pluripotent stem cells in most species (Takahashi and Yamanaka, 2006). As a starting point, we used bat embryonic fibroblast (BEF) cells isolated from wild-caught greater horseshoe bats (*Rhinolophus ferrumequinum*). Strikingly, the standard protocol (Hochedlinger and Jaenisch, 2015) that is highly effective in mice, and after adjustments in humans and other mammalian species (e.g., domestic dog *(Canis familiaris)*, domestic pig (*Sus scrofa)*, common marmoset (*Callithrix jacchus*), failed in bats. Even though the standard reprogramming protocol failed, it provided us with a crucial insight: the Yamanaka factors triggered the formation of rudimentary stem cell-like colonies even though they ceased to expand. This observation prompted us to suspect that the core pluripotency network might be conserved in bats, whereas the signaling cascades that usually shield this network from differentiation cues may differ. We therefore empirically altered the ratios and amounts of the reprogramming factors and, through a combinatorial approach, activated and blocked various cellular signaling pathways to ascertain if they enabled stem cell reprogramming in bats. To this end, we identified that a specific ratio of reprogramming factors, and the addition of Lif, Scf, the Pka activator forskolin, and Fgf2 to the culture medium, allowed for the uninterrupted growth of bat pluripotent stem cells (Figure 1A, detailed in the Materials and Methods section). Under these adjusted conditions, bat stem cell colonies typically appeared after 14-16 days of culture. These initial stem cell colonies were, however, not readily pickable and expandable using conventional EDTA (Versene)-, collagenase- or trypsin-based methods that are normally used to passage pluripotent stem cells from other species. The only effective method seemed to be lightly flushing the cells off the feeder cell layer after gentle treatment with low concentrations of EDTA.

### Bat Pluripotent Stem Cells

Bat iPSC colonies appeared tight and homogeneous. The cells had a large, apparent nucleus with one or two prominent nucleoli (Figure 1B, Figure S1A), and were filled with tiny vesicles not seen in other mammalian pluripotent stem cells (Figures S1A-E). Bat iPSCs expressed the pluripotency factor Oct4 as demonstrated by immunostaining (Figure 1C), and their proliferation rate was similar to that of human pluripotent cells. The cells retained a normal karyotype, with most cells containing 56 chromosomes (Figure S1F) and replicated in the absence of the exogenous reprogramming factors (Figure S1G) for now more than 100 passages without a change in morphology. RNA-seq analyses (at passage 22) revealed induced endogenous expression of canonical pluripotency-associated genes such as Oct4, Sox2 and Nanog (Figure 1D, Figure S2A, Table S1A). However, closer data inspection revealed that the expression profile did not fully match a single known pluripotency state. Instead, we saw factors indicative of the *naive* pluripotent state (e.g., Klf4, Klf17, Essrb, Tfcp2l1, Tfe3, Dppa, and Dusp6), expressed alongside genes typically found in the more advanced *primed* pluripotent cells (e.g., Otx2, Zic2), a phenomenon previously described for human cells under certain culture conditions (Cornacchia et al., 2019). Indeed, double immunostainings detecting four of the most commonly used primed/naïve factors, Otx2/Tfe3 and Tfcp2l1/Zic2, respectively, show co-expression of naïve and primed markers in most cells (Figure S2B). In contrast, germ cell factors such as Dnmt3l and Dazl were absent. Thus, while cellular heterogeneity might be at play, their uniform appearance makes it most likely that bat stem cells occupy a novel, yet-to-be-characterized pluripotent default state.

Next, we checked the effects of our reprogramming approach on the bat chromatin and epigenetic structures. A global epigenetic landscape survey using the assay for transposase-accessible chromatin with sequencing (ATAC-seq) revealed substantial chromatin configuration changes when bat fibroblasts transitioned into the pluripotent state (Figure 1E). Similarly, mapping the DNA methylome by reduced-representation bisulfite sequencing (RRBS) exposed major CpG methylation changes across the genome (Figure 1F, Figure S3A, Tables S1C, and S1D) after reprogramming. Finally, ChIP-seq for histone marks associated with active (H3K4me_3_) and developmentally repressed genes (H3K27me_3_) showed many changes (Figure 1G, Table S1E) and were also associated with pluripotency genes (Figure S2A). Approximately 18.2% of the bat stem cell genes were associated with a “bivalent” domain (H3K4me_3_ and H3K27me_3_; Figure 1H, Table S1E), a pluripotency chromatin hallmark initially found in human and mouse pluripotent cells (Bernstein et al., 2006). Interestingly, while there was overlap between human (Court and Arnaud, 2017) and bat bivalency genes there were also some bat- or human-specific genes (Figure S3B). Generally, there were strict correlations between an increase in gene expression and newly opened sites showing increased ATAC-seq signals and H3K4 trimethylation along with decreased H3K27 trimethylation and DNA methylation in their promoters (Figure S3C-E). Conversely, closed regions and gene shutdowns during the reprogramming process also corresponded to the absence of activating and presence of histone modifications, respectively (Figure 1I). However, there are instances when we see transcription and simultaneously active and repressive epigenetic marks, most likely as a result of spontaneous differentiation in our cultures (Figure S2A). Collectively, our results establish that our BiPS cells are reprogrammed both transcriptionally and epigenetically.

The transcriptional and epigenetic changes mentioned above suggest but are not definitive proof of developmental pluripotency. To obtain functional evidence, we subjected the bat stem cells to protocols optimized for directed differentiation into ectodermal, mesodermal, and endodermal fates (Figure 2A, Figure S4A). In each case, the cells responded to the altered culture conditions by shifting their morphology profoundly and turned positive for Pax6 (a marker for ectoderm), T (mesoderm), or AFP (endoderm), respectively. Since the cells used in this experiment were at an advanced passage (passage 37, an equivalent of about 6 months of continuous culture), the results also suggest that pluripotency can be maintained long-term. We next probed the developmental plasticity of bat stem cells by subjecting them to embryoid body (EB) differentiation, another classical *in vitro* pluripotency assay (Wiles and Keller, 1991). Again, the BiPS cells (referring to iPS cells derived from *Rhinolophus ferrumequinum*) differentiated and formed spherical arrangements typical for EBs (Figure 4B) that subsequently matured into elaborate three-dimensional structures positive for all three germ layer markers (Figure 2B). RNA-seq analyses of RNA isolated from the cells following monolayer differentiation and EB formation confirmed the respective cell fate changes (Figure 2C, Figure S4C-D, Table S2).

**Figure 2.**
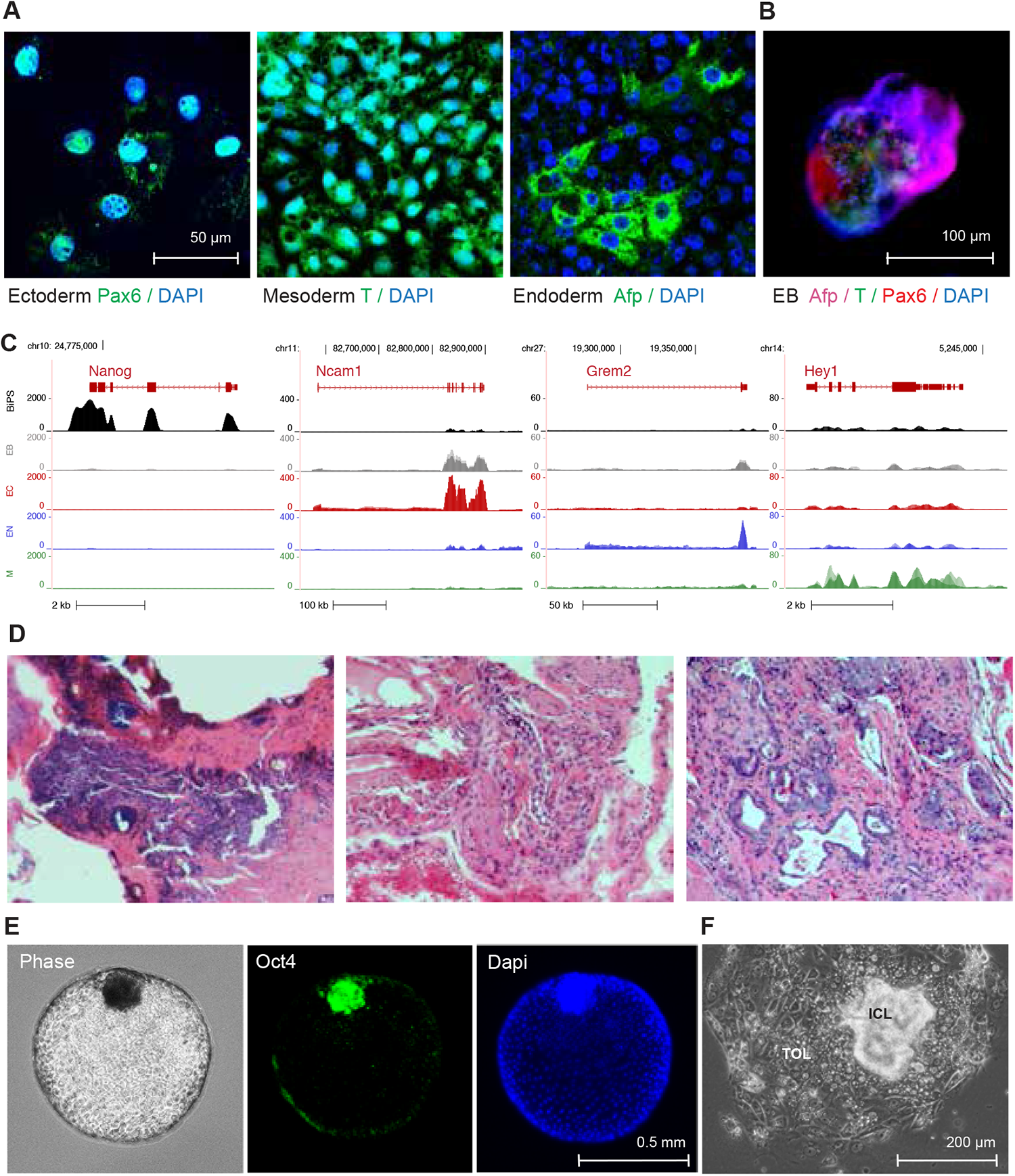
Differentiation potential of bat pluripotent stem cells. **(A)** Immunofluorescence microscopy images after staining with antibodies detecting the expression of lineage-specific markers Pax6, Afp or Brachyury (T) following specific directed differentiation into ectoderm, endoderm, or mesoderm, respectively. **(B)** Immunofluorescence images of embryonic bodies (EB) that formed after 3D-differentiation of BiPS cells and were stained with antibodies to detect markers specific to all three germ layers as in (A). **(C)** RNA-seq signals of selected lineage-specific marker genes in BiPS cells that underwent monolayer differentiation as in (A) or embryonic body differentiation as in (B). EB, embryonic body differentiation, EC, human ectoderm differentiation protocol; EN, human endoderm differentiation protocol; M, human mesoderm differentiation protocol. **(D),** Microscopic images of Hematoxylin-Eosin-stained sections of tumor tissue after injection of BiPS cells into immunocompromised mice exhibiting ectodermal (left), mesodermal (middle) and endodermal (right) features. **(E)** Images of floating blastoids that were obtained from BiPS cells after exposure to Bmp4 to capture their morphology by phase-contrast microscopy (left) and to detect Oct4 expression in inner-cell mass-like cell clusters after immunofluorescence staining (middle, right). **(F)** Phase-contrast microscopy image of a typical blastocyst outgrowth-like cell cluster that formed after attachment of blastoids to the cell culture vessel surface during Bmp4-induced differentiation as in (E). ICL, Inner cell mass-like; TLO, trophoblast-like outgrowth.

Next, we injected the BiPS cells into immunocompromised mice, because pluripotent stem cells typically form a particular tumor (teratoma) at the injection site (Damjanov and Andrews, 2007) that is composed of ectodermal, mesodermal, and endodermal cells arranged in a semi-chaotic fashion. While it took substantially longer than usual for pluripotent cells derived from other species, BiPS cells eventually formed a similar type of tumor after four to five months, albeit infrequently (33%) and relatively small (2-4 mm). The tumors consisted of immature tissue with epithelial, neural, and stromal characteristics (Figure 2D). Transcriptional profiling of pivotal genes previously reported critical for teratoma formation (Figure S6) revealed that while some genes are downregulated in bat iPSCs in comparison with mouse iPSCs (like Eras), other genes like the hyaluronidases (HAS) and ADP ribosylation factors (ARFs) are indistinguishable between the experimental groups, making it likely that the anti-tumor effect seen in the rudimentary teratomas is a complex phenomenon. While the host mice were severely immunocompromised and we had no access to immune-related tissues, the immaturity and delay in growth may suggest a yet to be characterized anti-tumorigenic property of bat stem cells similar to, for instance, naked mole rats (*Heterocephalus glaber)* (Miyawaki et al., 2016; Tan et al., 2017), which could also underlie the extended health span and cancer resistance reported in many bats (Munshi-South and Wilkinson, 2010; Wilkinson and Adams, 2019). Finally, we created embryo-like structures from bat stem cells using a modified blastoid protocol (Yu et al., 2021) (Figure 2E). These bat blastoids recapitulated critical aspects of preimplantation embryos, including an Oct4-positive inner cell mass, the cystic cavity, and a bilayered epithelium consisting of trophoblast and yolk sac cells. Replating these embryo structures resulted in their attachment with a flattened trophoblastic epithelium outgrowth and an expansion of the inner cell mass (Figure 2F). Our differentiation studies exemplify the unique potential of pluripotent bat stem cells to recapitulate important developmental events and serve as a powerful model to study the unique physiological adaptations of bats, including their reduced cancer phenotype. Finally, to see if our protocol is broadly applicable to bats, we created primary *Myotis myotis* fibroblast cells from 3 mm uropatagium (tail) biopsies of wild-caught adult bats. These fibroblasts were readily reprogrammable using our new “batified” Yamanaka protocol and yielded similar bat iPSCs that were Oct4 positive in immunostaining and differentiated into all three germ layers (Figure S5), suggesting that our protocol is applicable across the deepest basal divergencies in bats.

### Comparative Transcriptomics

Next, we investigated whether our new stem cell model can be used to gain novel insights into the unique evolutionary adaptations of bats. Phenotypic differences among species can be driven by evolutionary changes in gene expression (Romero et al., 2012). Therefore, given the unique adaptations of bats, we should detect bat-specific gene expression patterns in bat stem cells, allowing us to compare the transcriptomic ground state across species, which is one of the earliest ontogenic comparisons possible. We collected transcriptome profiles of pluripotent stem cells from phylogenetically divergent mammal species (*Mus musculus* (mouse), *Homo sapiens* (human), *Canis familiaris* (dog), *Sus scrofa* (pig), *Callithrix jacchus* (marmoset)) to compare them to our greater horseshoe bat (*Rhinolophus ferrumequinum*) data. Principal component analyses were performed to obtain a high-level overview of the number of commonalities and differences between bats and other mammals (Figure 3A). Remarkably, and supportive of our suspicion that bats are unique, all other mammals grouped together, in the PCA plot, while our bat stem cells formed a separate distinctive group, despite including other related laurasiatherian mammals (Jebb *et al*., 2020).

**Figure 3.**
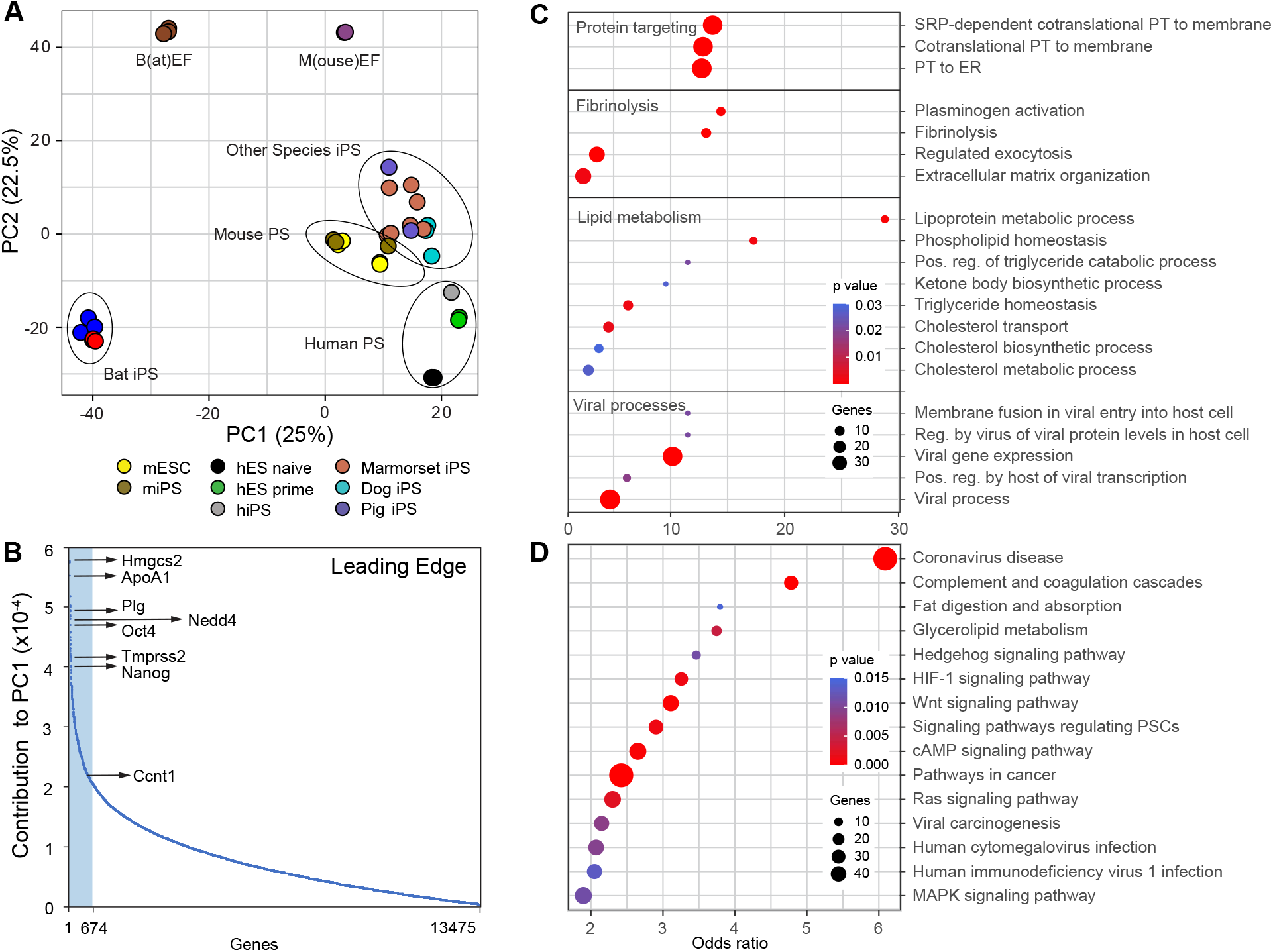
Distinct characteristics of bat pluripotent stem cells. **(A)** Principal component analysis of induced pluripotent bat stem cells (BiPS) in comparison to those derived from other species. h, human; m, mouse. PS, pluripotent stem cells, iPS, induced pluripotent stem cells, ES, embryonic stem cells, EF, embryonic fibroblasts. **(B)** Plot of genes that contribute to the differences of pluripotent bat and mouse stem cells as part of principal component 1 (PC1). Highlighted in light blue is the “leading edge” comprised of the top 5% of PC1-contributing genes. **(C)** Selected GO and **(D)** KEGG pathways identified to be significantly enriched among the top 5 % of PC1-contributing genes/leading edge genes defined in (B) were plotted by their odds ratio, with the color of each circle indicating the enrichment p-value and the size indicating the number of genes present in the respective category (see Data Table S3B and S3C for a full list of enriched gene sets). ER, endoplasmic reticulum; PT, protein targeting; Pos, positive; Reg, regulation.

We then determined the gene signature that contributed the most to the bat-specific gene expression profile. Specifically, we extracted the “leading edge,” corresponding to the top 5% of the genes that fortified the difference in principal component 1 (Figure 3B, Table S3A) when comparing bat with mouse pluripotent stem cells. The list covered genes belonging to a broad spectrum of transcription factors, kinases, metabolic and homeostatic enzymes. For instance, it included Hmg-CoA synthase Hmgcs2, apolipoprotein Apoa1, cyclin Ccnt1, plasminogen PLG, pluripotency factors Oct4 and Nanog, Tmprss2 which is required for SARS-CoV-2 entry in humans and the ubiquitin ligase Nedd4 among many other genes. Given the broad spectrum of categories we next asked if the leading-edge genes were enriched for any particular biological pathway in gene ontology analyses (Figure 3C, Table S3B). We expected the genes to primarily encode developmental controllers and indeed, some of the genes belonged to this class, but the vast majority fell into rather unexpected categories. Among the enriched functional families were proteins targeting membranes, including the endoplasmic reticulum, lipid and cholesterol biosynthesis, and fibrinogen production. However, the most prominent groups were viral gene expression, viral transcription, and many sets activated or suppressed after viral infection (Figure 3C, Table S3B). “Coronavirus disease” was by far the most significantly enriched category in any KEGG pathway (Figure 3D, Figure S7A, Table S3C). These results suggest that bat stem cells execute a program that in other mammalian cells is activated only after virus infection.

Interestingly, out of the set of leading-edge genes, only a total of eight genes showed significant evidence of positive selection in *R. ferrumequinum* (Figure S7B, Table S3D and S3E). Two of these genes, Col3a1, and Muc1, have roles in collagen formation in connective tissues (Kuivaniemi and Tromp, 2019), protect against pathogen infections (Kuivaniemi and Tromp, 2019) and showed evidence of selection in another bat species suggesting unique, bat-specific adaptations in these genes. Our results might indicate that the unique bat signature is likely the consequence of the presence of viral sequences triggering the expression of antiviral cellular programs and that most of the coding leading edge genes are not under positive selection pressure.

### Endogenous viruses in bat stem cells

Throughout their evolution, bats have absorbed diverse viral sequences into their genomes (Pourrut *et al*., 2009; Skirmuntt and Katzourakis, 2019). This is in line with our findings that multiple virus-infection-related gene categories are highly enriched in bat stem cells (Figure 3, Figure S7A). Since endogenized viral sequences are often awakened in the developmental tabula rasa state of pluripotency in humans and mice (Grow *et al*., 2015), we hypothesized that our bat pluripotent stem cells would display a particularly rich set of expressed endogenized viral sequences and antigens compared to other mammals.

To put the hypothesis that a particularly broad array of endogenized viral sequences would be re-activated in bat pluripotent stem cells to test, we looked first at endogenous retroviruses, which are abundant and diverse in bat genomes (Hayward *et al*., 2018; Jebb *et al*., 2020; Skirmuntt and Katzourakis, 2019; Zhuo et al., 2013). As a starting point, we picked anchor points of retroviral sequences, and mapping our RNA-seq from bat iPSCs revealed the expression of a markedly diverse set of retroviral families in bat pluripotent stem cells when compared to fibroblasts (Figure 4A and Table S4A). We detected not only previously characterized full-length bat retroviruses (Figure S8A) but also novel ones (e.g., RFe-V-MD1) that are transcriptionally activated during reprogramming (Figure 4B, Table S4B). Importantly, chromatin in the vicinity of expressed endogenous retroviruses opened up epigenetically during reprogramming, confirming our suspicion that the reprogramming process was responsible for revealing the diverse set of ERV sequences in our bat stem cells (Figure S8B).

**Figure 4.**
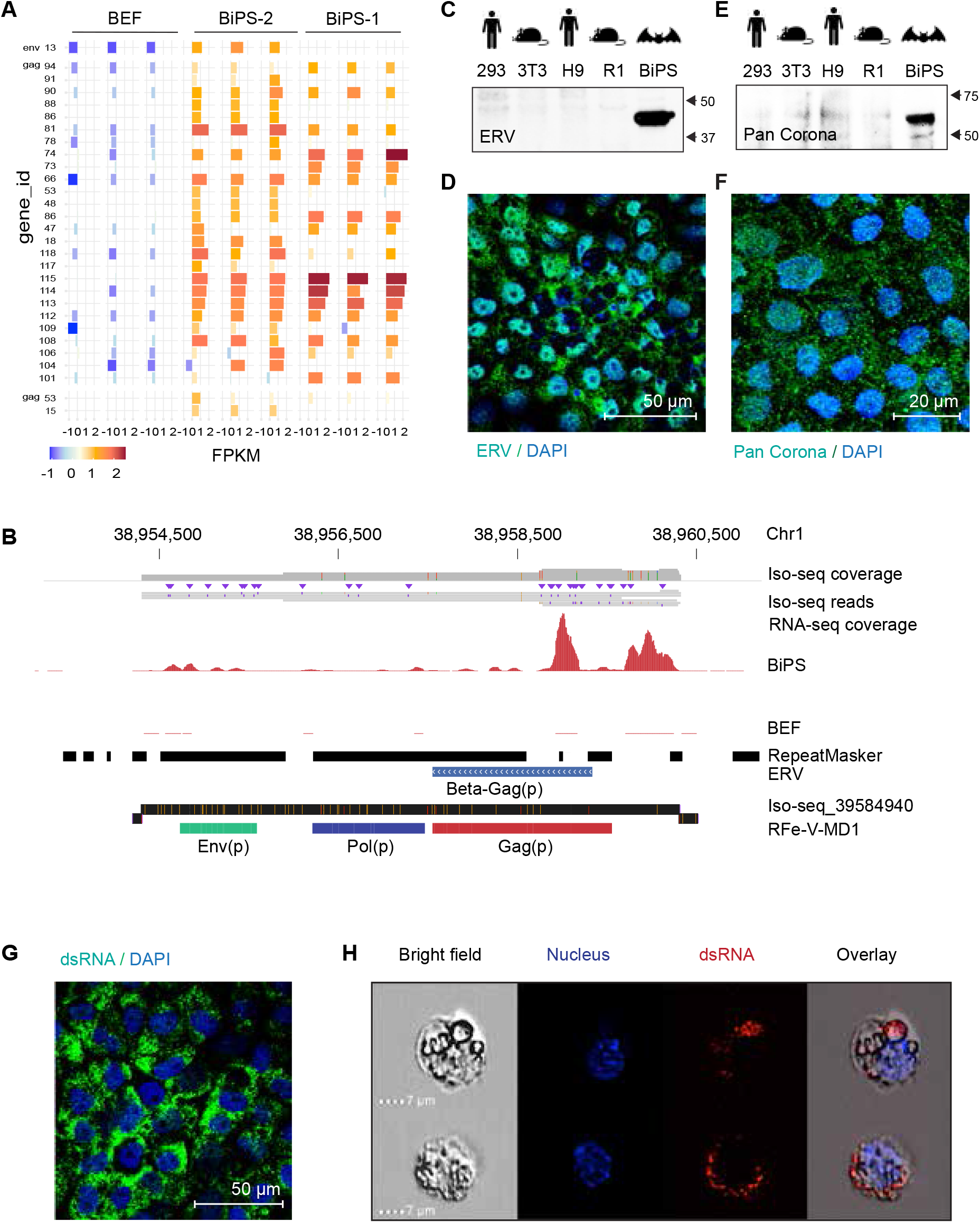
Reactivation of endogenized viral elements in bat pluripotent stem cells. **(A)** Expression of indicated ERV elements in bat embryonic fibroblasts (BEF) and iPS cells (BiPS) as determined by extracting the overlap between RNA-seq reads mapped to the *R. ferrumequinum* genome and known mapped ERV elements. Shown are the elements with the most evident differences (see Data Table S4 for a full list of expression data). **(B)** RNA and Iso-seq sequencing tracks for a newly discovered full-length retrovirus sequence, RFe-V-MD1, aligned to the *R. ferrumequinum* genome. The Iso-seq fragment represents a 6088 bp-long transcript **(C)** Western blotting in human 293FT (kidney tumor cells) and human embryonic stem cells (H9), mouse 3T3 (fibroblasts) and mouse embryonic stem cells (R1), and bat induced pluripotent stem cells (BiPS) with an ERV-specific antibody. **(D)** Immunofluorescence images of BiPS cells and bat embryonic fibroblast (BEF) cells after detection of the HERV K Cap protein (green); DAPI (blue). **(E)** Western blotting in human 293FT and H9, mouse 3T3 and R1, and BiPS cells with a pan coronavirus antibody known to be specific for the nucleocapsid; its reactivity includes but might not be limited to feline infectious peritonitis virus type 1 and 2, canine coronavirus (CCV), pig coronavirus transmissible gastroenteritis virus (TGEV) and ferret coronavirus. **(F)** Immunofluorescence images of BiPS cells after detection of the pan coronavirus antigen (green); DAPI (blue). **(G)** Immunofluorescence images of BiPS cells after detection of double-stranded RNA (Green) characteristic of RNA viruses; DAPI (blue). **(H)** ImageStream analysis after immunofluorescence staining of BiPS cells. A brightfield image, DyeCycle Violet nuclear staining (blue), dsRNA staining (red) and an overlay is shown for each representative cell.

To determine if the transcript-based findings indeed hold true at the protein level, we then determined whether ERV antigen was present in our bat stem cells. Indeed, western blotting and immunostaining revealed high levels of ERV antigen in bat stem cells that were not detected in human or mouse stem cells or fibroblasts (Figure 4C and 4D, Figures S8C and S8D). Additionally, our transmission electron microscopy (TEM) revealed electron-dense particles with lucent cores ranging in size from 50 to 100 nm that sometimes resembled previously reported viral-like particles (VLPs) (Figure S1D) (Grow *et al*., 2015). Although the exact nature of these particles needs to be explored further, virus-like activity was detectable in the supernatant (1.21 * 10^10^ viral particles per ml as determined in a retroviral assay and 0.3 ng/well in a direct reverse transcriptase assay). However, when we inoculated Vero cells with supernatant of our bat iPSCs in plaque assays, we did not detect any measurable cytotoxic effects in contrast to acute infectious virus particles that served as positive controls (Figure S8E). These findings imply that bat cells produce ERV antigen and, in some instances, possibly active endogenous viral-like assemblies at an unusual scale compared to other mammals.

In addition to ERVs, bat genomes have also assimilated a substantial number and diversity of endogenous viral elements (EVE) (Jebb *et al*., 2020). Hence, we attempted to obtain a broader portrait of integrated and expressed viral sequences. We developed new pipelines based on metagenomic classification of the stem cell RNA-seq data using either Kraken2 or Microsoft Research Premonition (Figure S9, Tables S5 andS 6), which included a series of strict classification (using the k-mer based Kraken2 and the alignment based Microsoft Premonition) and curation steps (*de novo* assembly of putative viral contigs and genome mapping) to identify true viral reads (Supplementary Material and Methods, Figures S10A and S10B). The analyses revealed that bat pluripotent stem cells display a variety of virus-associated endogenized sequences. For instance, Blast analysis of selected suspected viral Iso-seq reads as identified by our metagenomics method showed an unexpected region in the first intron of the Xpa gene (DNA damage and repair factor) on chromosome 12. The region showed homology to two human herpesvirus 4 isolates (HKD40 and HKNPC60), the human respiratory syncytial virus (Kilifi isolate), and a fragment of about 500 bp that was identified at the end of a SARS-CoV2 isolate from an infected patient (Figure S10C and 10D, and Data Table S7A). We also found a nearly 50% identical sequence to either *Scotophilus* bat coronavirus 512 - or *RaTG13* coronavirus (one of the bat coronaviruses most closely resembling SARS-CoV-2) covering most of the spike encoding sequences (Table S7A-C). Phylogenetic analysis revealed that these genomic sequences resembled the spike encoding genomic portion of viruses such as the human coronavirus 229E and human coronavirus OC43, respectively (Figure S10E). Interestingly, these regions are flanked by LINE-1 sequences (Figure S10F). This suggests the possibility that LINE elements are directly involved in the homing of viral RNA a possibility that was also explored recently in the context of SARS-CoV-2 (Zhang et al., 2021). Systematically scanning for expressed viral elements in our RNA we then assembled contigs and aligned them to viral and mammalian genomes using BLAST (Figure S9, Tables S6 and S7, and Supplementary Material and Methods, Data Blast A-P) using highly stringent parameters (Supplemental Material and Methods, Figure S11A). This procedure allowed us to identify previously unknown viral sequences and integration sites (Tables S6 and S7). While many of the viral alignments were difficult to distinguish from cellular genes (e.g., oncogenes) especially in pipelines that do not include genome mapping steps, we detected what appeared to be bona fide purely viral-like elements that were integrated in the bat genome and induced in our pluripotent stem cells.

We identified numerous predicted and new retroviral integration sites with homologies to the Koala retrovirus, Mason-Pfizer monkey virus, Jaagsiekte sheep retrovirus or Ovine enzootic nasal tumour virus, to name a few (Table S6 and S7; Figures S11A-C). Translations of the extended regions covered by mapped RNA-seq reads using *BLASTX* revealed similarities to the Jaagsiekte sheep reverse transcriptase and other known retroviral gag, pro and pol protein (Tables S7A, S7D, S7e, and S7F). An example of an integrated DNA virus sequence is a region within scaffold_m29_p_1 of the *R. ferrumequinum* genome which shows homologies to the Volepox, Variola, Squirrelpox and Monkeypox viruses (Figure S11D). Here, translation of the extended region with mapped RNA-seq reads uncovered homologies with the cowpox protein CPXV051 and the monkeypox C10L protein (Tables S7A and S7G). Another region that is worth noting is located within scaffold_m29_p_20. We first identified this region through a short sequence homology with the *Severe acute respiratory syndrome*-related coronavirus isolate Rs7907 coding for an N-terminal fragment of a dsRNA binding protein (Table S6, and Figure S11E). When extending this region to cover all mapped RNA-seq reads in the vicinity, a blast search identified several longer homologies with the White spot syndrome virus (Figure S11F, and Tables S7A and S7H). Interestingly, we find mapped RNA-seq reads in fibroblasts and iPS cells and that the genomic location borders a “gap” region in the genome, indicating that even more of this non-mammalian virus might be present in the bat genome. In summary, we conclude that, while confounding effects including genomic contaminations can affect the metagenomic classification process, it is highly likely that a sizable body of proviral sequences and sequence fragments inhabit BiPS cells.

To support our transcriptomics-based findings, we next looked for antigen markers linked with the RNA virus lifestyle because bats have traditionally shown an extreme affinity for RNA viruses and in some cases coronaviruses (Villarreal, 2009). We first stained our bat iPS cells with an antibody detecting a corona virus antigen. This was based on the facts that bats are known to host Corona viruses and early during the SARS-CoV-2 pandemic *Rhinopholidae* were discussed as host for SARS-CoV2, and our discovery of sequences that resemble Corona-viruses (Table S7, Figure S10). Indeed, we found the BiPS cells to be positive in immunofluorescence and western blot analyses when compared to fibroblasts and pluripotent stem cells from other species (Figures 4E and 4F, Figures S12A and S12B). Super-resolution microscopy showed clustered localization within the cytoplasm (Figure S12C). We then looked for the presence of double-stranded RNA in immunostaining which is thought to be a sign of replicative genomes from both positive-strand, double-stranded RNA and DNA viruses (Figure 4G). Super-resolution imaging (Figure S12D) showed that the dsRNA was present in micron-order-sized aggregates throughout the cytoplasm but essentially absent from the nucleus. Further, ImageStream analysis revealed an overlap between the dsRNA signal and the distinctive intracellular vesicles found in bat stem cells (Figure 4H, Figure S12E). However, the precise involvement of these vesicles in viral activities needs to be further investigated. Finally, in line with the pro-viral environment on transcriptional level, we found that bat stem cells infected with an exogenous Metapneumovirus (MPV) revealed a particularly permissive environment for viral persistence when compared with mouse stem cells, further underscoring the supportive nature of bat stem cells for viruses (Figure S13).

The results of our study provide proof-of-concept evidence that bat stem cells include a remarkable variety of sequences that are similar to viral genomic sequences. Additionally, our findings indicate that retroviruses and parts of endogenous viruses other than retroviruses are produced and active on a scale that is not generally seen in tumor or stem cell lines that originate from other animals or humans. We conclude that the transcriptionally permissive state of pluripotency can be exploited to discover novel bat viruses and derivative sequences that likely play an essential role in bat physiology and their ability to host viruses.

## Discussion

Bats have evolved an unusual lifestyle amongst mammals as they fly, use echolocation, and have a curious affinity for viruses. One possibility is that bats evolved a tolerance for viruses by evolving changes in their innate immunity resembling the virus evasion mechanisms of the mammalian immune response. Another possibility is that bats evolved mechanisms for a cellular program to support viral replication and persistence, comparable to how viruses manipulate the host cell. Our results support both perspectives.

Indeed, our results show that a potentially significant contingent of endogenous and exogenous viral products are present in bat pluripotent stem cells without severely compromising their ability to proliferate and grow and that this goes beyond how other pluripotent stem cells react to viruses. Viruses typically adapt their replication cycles to a particular cell type. Thus, one would not expect the pluripotent stem cell state to align with viruses’ often specialized requirements (Macfarlan *et al*., 2012). Nevertheless, our data suggest that in bats, the pluripotent state serves as an ‘umbrella’ host for a highly divergent viral contingent. We propose that our culture model can help to carefully dissect the necessary balance for tolerance of viral infections. Our new bat stem cell system will also provide insights into bats’ potential role as virus reservoirs and the relationship between bats and viruses. *In vitro* differentiation into immune cells and tissues, like lung or gut epithelium, will illuminate emerging viruses and how bats can tolerate viral infections, and in turn allow us to better prepare for future pandemics.

Bats are a critically needed new model organism to better understand disease tolerance, but limited access to animal and cell models has hindered their study (Rasweiler et al., 2009; Wang *et al*., 2021). Bat breeding colonies for some species are notoriously challenging to establish; most bat species are protected worldwide, and primary bat cell lines typically have a limited *in vitro* lifespan (Wang *et al*., 2021). Therefore, pluripotent stem cells offer a research tool that is a *sine qua non* for bat research. Once established, pluripotent stem cells divide indefinitely in culture and are well amenable to gene editing and molecular studies (Takahashi and Yamanaka, 2016). Most importantly, pluripotent stem cells retain the ability to differentiate into any cell type in the body and are often used as a springboard for sophisticated tissue culture models (Lancaster et al., 2013) and, more recently, virus studies (White et al., 2021; Yang et al., 2020). Future research on bat stem cells will directly impact every aspect of understanding bat biology, including bats’ amazing adaptations of flight, echolocation, extreme longevity, and unique immunity. While bat genomes are the natural starting point for studying such adaptations and are being generated by the Bat1K consortium (Teeling *et al*., 2012; Teeling *et al*., 2018), our pluripotent stem cell system will enable specific bat tissue studies, and organoids will let us test more complex relationships, gene editing, and specific evolution hypotheses especially since our bat stem cell conditions do not support reprogramming or stem cell self-renewal in mice. Bat stem cell lines and differentiated progeny will help address tantalizing physiology questions, provide the required tools for validation, and utilize the genomic basis of rare adaptations found in bats.

While metagenomic classification of only recently assembled genomes and transcriptomes has limitations in the context of diverse mammalian genomes and will need to be verified using future curated assemblies and at the protein level, it is a solid starting point to carefully revisit the plethora of genomic and expressed bat viral sequences and to test if they are integrated sequences to defend against viruses and microbes and encode viral proteins in a self-vaccination scheme or are near full-length viruses to manipulate host physiology. Also, the reactivation of endogenized virus fragments exposes only part of the host tissue response and needs to be investigated in other cell types and acute viruses. Furthermore, how closely the new pluripotent stem cell lines resemble different inner cell mass (ICM) states will only be addressable by comparing them to bat ICM cells and generation of bat chimeras and deriving pluripotent stem cell lines from other bats besides horseshoe bats, which becomes feasible with our revised Yamanaka protocol. This current proof-of-concept study establishes bat stem cells in particular and bats in general, as tantalizing novel model systems, allowing us to both elucidate the diversity of viruses that bats can survive and the molecular adaptations that enable bats asymptomatically tolerate these viruses, thus providing new insights into both disease surveillance and future therapeutics.

## Supporting information

BLAST files

## Acknowledgements

We thank Prof. Eloy Revilla EBD-CSIC’s Director, the EBD-CSIC’s Bio-Ethical Committee, the CABIMER, the ‘Consejería de Medio-Ambiente’ of the ‘Junta de Andalucía’ and the ‘Subdirección General de Acuerdos Sanitarios y Control en Frontera of the Spanish Ministery of ‘Agricultura, Pesca y Alimentación for their support and help to obtain the export permits in a record time and many other people for making the shipment and delivery of the samples from Seville, Spain to New York, USA under the COVID-19 pandemic’s most arduous conditions possible. We also thank Michael Schotsaert and Carles Martinez for helping with the bat tissue import, Allison Sova and Bill Williams from the Microscopy Core and Advanced Bioimaging Center at the Icahn School of Medicine for preparing the cells for the electron microscopy and imaging, as well as Glenn Doherty and Nikolas Tzavaras for their assistance with the confocal microscopy. We also thank the Genomics and the Biorepository and Pathology Core Facility members at the Icahn School of Medicine at Mount Sinai for performing the RNA-seq and histology, respectively. This work was supported in part through the computational resources and staff expertise provided by Scientific Computing at the Icahn School of Medicine at Mount Sinai. ECT is supported by Irish Research Council Laureate Award IRCLA/2017/58 and Science Foundation Ireland Future Frontiers 19/FFP/6790. GMH is funded by a University College Dublin Ad Astra Fellowship. MD and TPZ are supported by the Huffington Foundation. AGS is supported by HR0011-19-2-0020, awarded by DARPA and Grant No. W81XWH-20-1-0270, awarded by Department of Defense (DoD). This work was also partially supported by NIAID grant U19AI135972 and by CRIPT (Center for Research on Influenza Pathogenesis and Response), a NIAID supported Center of Excellence for Influenza Research and Response (CEIRR, contract #75N93019R00028) to AGS and by grant 2021-244135(5384) from the Open Philanthropy Project Fund to AGS and TPZ. JJ and CI are supported by the Spanish Ministerio de Ciencia e Innovación (SAF2017-89355-P). AM and MH are supported by the LOEWE-Centre for Translational Biodiversity Genomics (TBG).

## Data Availability

Data generated during this study have been deposited in the Gene Expression Omnibus (GEO) and the accession code will be made available.

## Code Availability

All software packages and their accessibility are described in the Material and Methods sections.

## Biological Material Availability

The BiPS cell lines are available from the Zwaka Lab upon request.

## Additional Information and Files

Supplementary Information including materials and methods with additional references.

Figures S1-13

Data Tables S1-7

Data BLAST SA-P (please note that the file names correspond to the steps indicated in Figure S9 and Table S7A-P).

## Supplemental Information

### Note

Careful cataloging of acute exogenous bat viruses, tissue infection, persistent viruses, and endogenized viruses in geographically relevant regions (Wacharapluesadee et al., 2021; Zhou et al., 2021) will reveal novel members of diverse retroviral and non-retroviral sequences, potentially impacting host, and novel emerging viruses. It will also uncover new rationales for virus persistence, including immune-modulatory strategies (Wang et al., 2011), symbiotic protection against other pathogens (Barton et al., 2007; Roossinck M.J., 2011; Eaton et al., 2006), biological warfare that bats use to deploy viruses (Wang et al., 2011), novel mammalian adaptive piRNA or CRISPR-like systems Ophinni et al., 2019) and the augmentation of evolutionary processes (Feschotte and Gilbert, 2012). While the events we specifically study in pluripotent stem cells do not directly impact those occurring in specific adult cells we propose that pluripotent stem cells, in their own right, are an important cell system, when studying native immunity and viruses. They share a highly conserved (Endo et al., 2020) and immunologically relevant common genetic program with somatic cells (Wu et al., 2018). Pluripotent stem cells in the embryo and the related trophoblastic tissue must establish their immunological barrier against the maternal tissue and are subject to viral infection, as also noted in bats (Aikawa et al., 2014). Furthermore, as some fundamental cell biological properties might be shared between stem cells (Ivanova et al., 2002; Ramalho-Santos eet al., 2002), the finding may also extend to other stem cells often at risk of infection like HSCs (Carter et al., 2011), NSCs (Li et al., 2016; Janssens et al., 2018), and early human embryos where for instance rubella virus can destroy the conceptus or cause severe congenital defects (Naeye and Blanc, 1965). Also, most basic native immunity systems with pathogen pattern sensing and inflammatory responses, including inflammasome, NFKB, and interferon are basic tools largely present in their most ancient ancestors and broadly shared between species and cell types (Daugherty et al., 2012; Siddle and Quintana-Murci, 2014). Finally, viral RNA products present in bat cells might also represent a potential source for RNA recombination upon infection with exogenous homologous viruses, which could be a driver for viral evolution in bats.

Our new bat stem cell system may also provide a much-needed experimental substrate to parse the tantalizing hypothesis that viruses and hosts are more entangled and that viruses are fully competent agents and editors of host biology (Witzany et al., 2010). Therefore, viruses must be rich sources of new evolutionary instructions, especially given their extreme genetic adaptability and ability to transition between the living and chemical worlds (Villarreal et al., 2000; Villarreal L.P., 2007). Allowing viral evolutionary processes to unfold in our bat cell lines and mapping out changes to the host transcriptome will be the first steps toward understanding host-virus editing interactions.

### Methods Details

#### Bat embryonic fibroblast isolation

An embryo (approximately developmental stage 20) acquired from a Spanish *Rhinolophus ferrumequinum* bat was cut into several pieces while removing the head and as much of the inner organ tissue as possible. The pieces were then flushed with PBS and processed separately. The tissue was covered with 0.05% trypsin, minced with a scalpel, and incubated in a cell culture incubator at 37°C and 5% CO_2_ for 45 minutes. The trypsin was deactivated with fibroblast medium consisting of DMEM (Life Technologies, 10569-010), 10% fetal bovine serum (Sigma, F4135), 0.1 mM MEM Non-essential amino acids (Life Technologies, 11140-050), 2 mM GlutaMax supplement (Life Technologies, 35050-061) and Penicillin-Streptomycin (10 U/ml and 10 µg/ml, respectively; Life Technologies, 15140122). The cells were broken up by pipetting up and down 20 times, collected by centrifugation, transferred to a gelatin-coated (Sigma-Aldrich, G1890) T75 cell culture-treated flasks (Falcon, 353136) in 15 ml of fibroblast medium, and cultured at 37°C and 5% CO_2_. After 3 days, when reaching ∼80% confluency, the attached cells were washed with DPBS (Life Technologies, 14190144), treated with 0.05% trypsin-EDTA (Life Technologies, 25300-120) to obtain a single cell solution and either split at a ratio of 1:4 or used directly in a reprogramming experiment.

#### *Myotis myotis* sampling and isolation of fibroblasts from tail biopsies

M. *myotis* were sampled in Morbihan, Brittany in North-West France, July 2021 in accordance with the permits and ethical guidelines issued by ‘Arrêté’ by the Préfet du Morbihan and the University College Dublin ethics committee. This population has been transponded and followed since 2010 as part of on-going mark-recapture studies by Bretagne Vivante and the Teeling laboratory (Huang et al., 2019). Once captured, all bats were placed in individual cloth bags before processing. A single 3 mm biopsy was taken from the outstretched uropatagium of each bat using a sterile biopsy punch and immediately submerged in a Cryotube (Sartedt, 72.379) with 2ml of DMEM cell culture medium (Gibco, 11995-065) supplemented with 20% FBS (Gibco, 10500064), 1% NEAA (Gibco, 11140068) and 1% Antibiotic-Antimycotic containing Streptomycin, Amphotericin B and Penicillin (Gibco, 15240096), maintaining as sterile conditions as possible. All bats were offered food and water and rapidly released after processing. Biopsies were then stored at 4℃ and transported to the laboratory for processing within 6 days. Samples were further processed through a cell extraction methodology similar to a previously established protocol (Kacprzyk et al., 2021) with a few modifications. The samples were rinsed with DPBS (Biowest, L0615-500) and cut finely within a minimal amount of cell culture medium using sterile blades (Swann-Morton, 0208) to result in six 0.5 mm pieces. These pieces were then transferred aseptically to a cryotube containing cell culture medium and incubated for 18 hours with collagenase type II at 37°C with 5% CO2 to allow for digestion. The pieces were collected by centrifugation for 5 minutes at 300 rcf, resuspended in 2 ml of fresh cell culture medium and transferred to a 35 mm cell culture treated plate (Corning, CLS430165) for initial P1 expansion. Cell were then fed every 2-3 days with cell culture medium as stated but a reduced 0.2% concentration of Antibiotic-Antimycotic. For the first feeding a ⅔ media change is performed to avoid sudden changes in antibiotic-antimycotic concentration from 1% to 0.2%. When the cells reached 70% confluency, they were transferred to a T25 (Cellstar, 690175) in cell culture medium after treatment with 0.05% Trypsin (Gibco, 25300054) and were fed every 2-3 days as necessary. At 85% confluency, the cells were trypsinized as before and 1×10^6 cells were frozen in 1 ml cell culture medium containing 10% DMSO (Sigma, D8418-100ML).

#### Reprogramming of bat and mouse fibroblasts

150,000 embryonic *Rhinolophus ferrumequinum* fibroblasts at passage 2, adult *Myotis myotis* at passage 3 or CF1 mouse embryonic fibroblasts at passage 3 were resuspended in 1 ml of fibroblast medium and mixed with Sendai-virus particles containing the reprogramming factors Oct4, Sox2, cMyc and Klf4 (CytoTune iPS 2.0, Invitrogen, A16517) with a final multiplicity of infection (MOI) of 10, 10, 10, 20, respectively. The cells were plated on one gelatin-coated well of a 6-well plate (10 cm^2^, Corning, 353046) and cultured at 37°C with 5% CO_2_. The medium was replaced every 24 hours. 6 days after transduction, the cells of each well were collected by treatment with 0.05% trypsin-EDTA, seeded at a density of 50,000 cells per 60 cm^2^ on irradiated CF1 mouse embryonic fibroblasts (MEFs; Gibco, A34180) in fibroblast medium. After 24 hours, the medium was switched to 50 % fibroblast medium and 50% pluripotent stem cell (PSC) medium consisting of DMEM/F-12 (Life Technologies, 11330-057), 20% knockout serum replacement (Life Technologies, 10828-028), 0.1 mM MEM Non-essential amino acids (Life Technologies, 11140-050), 2 mM GlutaMax supplement (Life Technologies, 35050-061), Penicillin-Streptomycin (10 U/ml and 10 µg/ml, respectively; Life Technologies, 15140122), 100 µM 2-mercaptoethanol (Fluka, 63689), and 40 ng/ml FGF2 (R&D Systems, 233-FB). From then on, the medium was replaced every day with PSC medium until day 14 when the FGF concentration was increased to 100 ng/ml and the medium was supplemented with 10^4 U/ml Leukemia inhibitory factor (Millipore, ESG1107), 100 ng/ml SCF (R&D Systems, PHC2111) and 20 nM Forskolin (Sigma, F6886). Colonies appeared 14 to 16 days after transduction, were picked on day 20 and expanded on irradiated MEFs with Gentle Cell Dissociation Reagent (GCDR, StemCell Technologies, 07174). After that, cells were passaged approximately every 5 days, or when they were confluent at a ratio of 1:6 to 1:12 on irradiated MEFs. Cell and colony morphology were recorded with an EVOS digital inverted microscope (Invitrogen).

#### Mouse ES cell culture

R1 mouse ES cells were cultured as described before (Dejosez et al., 2008).

#### Karyotyping

Cells were treated with 100 ng/ml Colcemid solution (Life Technologies, 15210040) for 16 hours, then treated with 0.05% trypsin-EDTA for 15 minutes and filtered through a 40 µm cell strainer to remove clumps. Cells were collected by centrifugation, resuspended in 1 ml 0.075 M potassium chloride (Sigma-Aldrich, P9327) and incubated for 20 minutes at room temperature. 0.5 ml fixative [1 part glacial acetic (Fisher Scientific, A38-212) mixed with 3 parts methanol (Sigma-Aldrich, A412-4)] were added, cells were collected as before, resuspended in 4 ml fixative, and incubated for 20 minutes at room temperature. The fixation step was repeated, the cells collected as before and all but about 200 µl of the fixative was removed. The cells were resuspended in the remaining fixative and dropped onto slides that were precooled at -20°C. The slides were airdried and the cells stained for 10 minutes with Giemsa Staining solution consisting of 1 part Giemsa solution (Life Technologies, 10092013), and 3 parts Gurr buffer (Invitrogen, 10582013). The slides were washed with water, dried, and mounted in Cytoseal 60 (Thermo Scientific, 23-244257). High-resolution pictures of chromosome spreads were acquired with an AxioObserver microscope (Zeiss) using the 100x oil objective.

#### RT-PCR

mRNA was extracted with the RNeasy Mini Kit (Qiagen, 74104). 500 ng of each sample were used to generate cDNA by reverse transcription using the SuperScript™ IV VILO™ Master Mix (Invitrogen, 11756050). 2 µl of the cDNA were used to detect the presence of Sendai virus transcripts using GoTaq Green Polymerase (Promega, M7123), and the oligos as recommended in the CytoTune iPS 2.0 kit (Invitrogen, A16517). Gapdh was amplified as loading control using oligos with the following sequence: Z25-132:GAPDH_F1_GHB: TGGTGAAGGTCGGAGTGAAC and Z25-133:GAPDH_R1_GHB: GAAGGGGTCATTGATGGCGA). The PCR products were analyzed on a 2% agarose gel containing ethidium bromide.

#### Immunofluorescence staining

Cells were plated on µ-slides (Ibidi, 80286). After 4 days, cells were washed once with DPBS and fixed with Cytofix/Cytoperm solution (Becton Dickinson, BDB554714) for 20 minutes at 4°C. Cells were rinsed with Perm/Wash buffer (Becton Dickinson, BDB554714) and then incubated overnight at 4°C in Perm/Wash buffer containing primary anti-Afp (R&D Systems, AF1369) anti-Pax6 (BioLegend, 901301), J2 anti-dsRNA (Scicons, 10010200), anti-(gag/pol)HERVK (Austrial Biological, HERM18315) or FIPV3-70 anti-Pan Corona (Life Technologies, MA1-82189) or directly conjugated anti-Oct3/4-AF488 (Santa Cruz, sc-5279-AF488), anti-Brachyury (R&D Systems, IC2085G), anti-Otx2 (R&D Systems, AF1979), anti-Zic2 (Abcam, ab150404), anti-Tfe3 (Sigma Aldrich, HPA023881) or anti-Tfcp2l1 (R&D Systems, AF5726) in a 1:50 (anti-Oct3/4) or 1:100 dilution (all others). Cells were rinsed and washed 3 times for 2 minutes with Perm/Wash solution at room temperature followed by a 1-hour incubation with a 1:200 dilution of the corresponding secondary antibodies (Donkey anti-chicken-Cy3, Millipore, AP194C; Goat anti-chicken-AF488, Life Technologies, A-11039; Donkey anti-rabbit-AF647, Life Technologies, A-31573; Goat anti-rabbit-AF488, Life Technologies, A-10034; Goat anti-mouse-AF488, Life Technologies, A-11029) in Perm/Wash buffer. Cells were rinsed, washed twice for 2 minutes with Perm/Wash Buffer and then incubated for 5 minutes with Perm/Wash buffer containing 2 drops per ml NucBlue Dapi stain (Invitrogen, R37606). The buffer was removed, and the cells were cover-slipped in Prolong Dimond antifade mounting medium (Invitrogen, P36965). Images were acquired with an AxioObserver fluorescence microscope with Apotome (Zeiss). For the simulated emission depletion (STED) microscopy (super-resolution), the cells were plated on coverslips that were placed in wells of 6-well plates. The staining was performed as described above but with a 1:200 dilution of the Abberior Star 635P (Abberior, ST635P) secondary antibody in Perm/Wash buffer. Cells were rinsed, washed twice for 2 minutes with Perm/Wash Buffer and then incubated for 5 minutes with Perm/Wash buffer containing 2 drops per ml DyeCycle Violet stain (Invitrogen, V35003). The coverslips were mounted face down on glass slides with Prolong Dimond antifade mounting medium (Invitrogen, P36965). Images were acquired with a TCS SP8 confocal microscope with STED 3x and White Light Laser (Leica) with a 100x oil objective. 405 nm and 594 nm lasers were used for excitation and 775 nm laser for depletion. Image resolution obtained was 19.8 µm by 19.8 µm using a zoom factor of 6x.

#### RNA-seq, differential expression analyses and visualization

For RNA-seq, RNA was extracted from BiPS cells at passage 22 and BEFs at passage 3. Total RNA was extracted with the RNeasy kit (Qiagen, 74104) following the manufacturer’s recommendations including the DNase (Qiagen, 79254) digest, and eluted in 50 µl RNase/DNase free H_2_O. RNA-seq libraries were prepared with the SMART-Seq v4 Ultra Low Input kit (Takara Bio, undifferentiated cells) or the Stranded Total RNA with Ribo-Zero Plus kit (Illumina, differentiated cells) and 100 bp paired-end sequencing reads (PE100) were generated by Illumina sequencing (NovaSeq 6000 S1) to a depth of at least 50 million reads (100 million total reads). The quality of the reads from the RNA sequencing was analysed with FastQC v0.11.9 (Andrews, 2010), and visualized using MultiQC v1.9 (Ewels et al., 2016). The mean phred score was around Q35 across each base position in the BEF and BiPS samples and no filtering or processing was necessary. In the differentiated samples (EB, endoderm, mesoderm, ectoderm differentiation), the quality of the first nucleotide was less than Q20 in many cases and the reads were processed with Trimmomatic v0.39 (Bolger et al., 2014). To carry out the differential expression analysis, the genome of *Rhinolophus ferrumequinum* was used as reference genome (RefSeq accession GCF_004115265.1 assembled and annotated by the Vertebrate Genomes Project (https://vertebrategenomesproject.org) or GenBank accession GCA_014108255.1 assembled and annotated by the Bat1K project (https://bat1k.com) as indicated. The reads were mapped with HISAT2 v2.2.1 (Kim et al., 2019), the .sam files resulting from each mapping were converted into .bam files and indexed using SAMtools v1.10 (Li et al., 2009) and the reads mapped against each gene were counted using featureCounts v2.0.1 (Liao et al., 2014). The differential expression analysis was performed with DESeq2 v1.10.1 (Love et al., 2014). To visualize the RNA-seq data in the UCSC genome browser, bigwig files were generated using the bamCoverage command from deepTools (https://deeptools.readthedocs.io/en/develop/content/tools/bamCoverage.html; Ram-irez et al., 2016).

#### MA Plot

The MA plots were generated based on the DESeq2 (see above) results with the ggmaplot function (https://rpkgs.datanovia.com/ggpubr/reference/ggmaplot.html) from the R package ggpubr (https://rpkgs.datanovia.com/ggpubr/). Genes are indicated by dots, plotted by their log2 fold change between bat fibroblast and pluripotent stem cells and the log2 mean of normalized counts (ratio of means). Blue dots indicate genes with an adjusted p value of (or FDR) of <0.05 and a fold change of 2 (log2 fold change of 1), red dots indicate genes with an adjusted p value (or FDR) of <0.05 and fold change of -2 (log2 fold change of -1). Dotted lines are drawn at fold change of 2/-2 (log2 fold change of 1/-1).

#### Heatmap

The heatmap of pluritest genes, with ClustVis (Beta), a web tool for visualizing clustering of multivariate data (Metsalu et al., 2015) https://biit.cs.ut.ee/clustvis/). Rows are centered; unit variance scaling is applied to rows. Both rows and columns are clustered using correlation distance and average linkage.

#### ATAC-seq

ATAC-seq and bioinformatics analysis to detect open chromatin in bat fibroblasts and bat pluripotent stem cells was performed by Active Motif from 100,000 cryopreserved cells (ATAC-seq service, 25079). In brief, nuclei were isolated, and libraries of open chromatin were prepared with the Nextera Library Prep Kit (Illumina) by Tn5 tagmentation. The tagmented DNA was purified using the MinElute PCR purification kit (Qiagen), amplified with 10 cycles of PCR, and purified using Agencourt AMPure SPRI beads (Beckman Coulter). 42 bp paired-end sequencing reads (PE42) were generated by Illumina sequencing (using NextSeq 500) to a depth of at least 83 million total reads and mapped to the GCA_004115265.2 genome (Ensembl, annotation version 102) using the BWA algorithm with default settings (“bwa mem”). Alignment information for each read was stored as BAM file. Only reads that passed the Illumina’s purity filter, aligned with no more than 2 mismatches, and mapped uniquely to the genome were used in the subsequent analysis. Duplicate reads (“PCR duplicates”) are removed. Genomic regions with high levels of transposition/tagging events were then determined using the MACS2 peak calling algorithm (Zhang et al., 2008). To identify the density of transposition events along the genome, the genome was divided into 32 bp bins and the number of fragments in each bin was determined. The data were then normalized by reducing the tag number of all samples by random sampling to the number of tags present in the smallest sample. Peak metrics between samples were compared by grouping overlapping Intervals into “Merged Regions”, which are defined by the start coordinate of the most upstream Interval and the end coordinate of the most downstream Interval (= union of overlapping Intervals; “merged peaks”). In locations where only one sample has an Interval, this Interval defines the Merged Region. Intervals and Merged Regions, their genomic locations along with their proximities to gene annotations and other genomic features were determined and average and peak (i.e. at “summit”) fragment densities were compiled. The sequencing tracks (number of fragments in each 32 bp bin stored as “.bigWig” file) were visualized with the UCSC genome browser.

#### Reduced representation bisulfite sequencing (RRBS)

Reduced representation bisulfite sequencing of bat fibroblasts and pluripotent stem cells was performed by Active Motif (RRBS Service, Active Motif, 25069). Briefly, 500,000 cells were provided as a frozen pellet. Genomic DNA was isolated, and 100 ng were digested with TaqaI (NEB R0149) at 65°C for 2 hours followed by MspI (NEB R0106) at 37°C overnight. Following enzymatic digestion, samples were used for library generation with the Ovation RRBS Methyl-Seq System (Tecan, 0353-32) following the manufacturer’s instructions. In brief, digested DNA was randomly ligated, and, following fragment end repair, bisulfite converted using the EpiTect Fast DNA Bisulfite Kit (Qiagen, 59824) following the Qiagen protocol. After conversion and clean-up, samples were amplified resuming the Ovation RRBS Methyl-Seq System protocol for library amplification and purification. 75 bp single-end sequencing reads (SE75) were generated by Illumina sequencing (using NextSeq 500) to a depth of at least 27 million reads (total of 54 million reads), with at least 2.9 million covered CpGs. The reads were mapped to the GCA_004115265.2 genome (Ensembl, annotation version 102) and the percentage of methylation at CpG sites across the genome was calculated. To visualize the methylation ratios aligned to the gnome with the UCSC genome browser, the methylation ratio files containing the methylation ratio for each chromosomal position were first converted to bed files, that were then used to generate bigwig files with the “bedGraphToBigWig v4 tool” (https://www.encodeproject.org/software/bedgraphtobigwig/). Correlation scatter plots were generated to show the level of methylation at common CpG sites.

#### Chromatin immunoprecipitation sequencing (ChIP-seq)

5 million cells were fixed cells in 1% formaldehyde by adding 1/10 volume of freshly prepared formaldehyde solution (11% formaldehyde, 0.1 M NaCl, 1 mM EDTA, pH 8.0, 50 mM HEPES, pH 7.9) to the existing medium. Cells were agitated for 15 minutes at room temperature and the fixation was stopped by addition of 1/20 volume of 2.5 M glycine solution (final concentration of 0.125 M) to the existing medium and incubation at room temperature for 5 minutes. The cells were scraped off the wells, collected by centrifugation at 800 g and washed with 10 ml chilled 0.5 % Igepal in PBS per tube by pipetting up and down. Cells were pelleted by centrifugation as before and resuspended in 10 ml chilled PBS-Igepal containing 1 mM PMSF. Cells were collected as before, and the cell pellet was snap-frozen in liquid nitrogen. Further processing, chromatin immunoprecipitation and bioinformatics analysis to detect H3K4me3 and H3K27me3 was performed by Active Motif (HistoPath ChIP-seq service, 25001). In brief, chromatin was isolated by adding lysis buffer, followed by disruption with a dounce homogenizer. Lysates were sonicated and the DNA sheared to an average length of 300-500 bp with an EpiShear probe sonicator (Active Motif, 53051). Genomic DNA (Input) was prepared by treating aliquots of chromatin with RNase, proteinase K and heat for de-crosslinking, followed by SPRI beads clean up (Beckman Coulter) and quantitation with Clariostar (BMG Labtech). An aliquot of chromatin (20 µg) was precleared with protein A agarose beads (Life Technologies). Genomic DNA regions of interest were isolated using 4 µg of antibody against H3K4me3 (Active Motif, 39159) or H3K27me3 (Active Motif, 39155). Complexes were washed, eluted from the beads with SDS buffer, and subjected to RNase and proteinase K treatment. Crosslinks were reversed by incubation overnight at 65°C, and ChIP DNA was purified by phenol-chloroform extraction and ethanol precipitation. Illumina sequencing libraries were generated from the ChIP and Input DNAs with the standard consecutive enzymatic steps of end-polishing, dA-addition, and adaptor ligation. After a final PCR amplification step, 75-nt single-end (SE75) sequence reads were generated by Illumina sequencing (using NextSeq 500) to a depth of at least 36 million reads per sample and mapped to the GCA_004115265.2 genome (Ensembl, annotation version 102) using the BWA algorithm with default settings. Duplicate reads were removed, and only uniquely mapped reads (mapping quality >= 25) were used for further analysis. Alignments were extended in silico at their 3’-ends to a length of 200 bp, which is the average genomic fragment length in the size-selected library and assigned to 32-nt bins along the genome. The resulting histograms (genomic “signal maps”) were stored in “bigwig” files. To find peaks, the generic term “Interval” was used to describe genomic regions with local enrichments in tag numbers. Intervals were defined by the chromosome number and a start and end coordinate. Peak locations were determined using the MACS algorithm (v2.1.0) with a cutoff of p-value = 1e-7 (Zhang et al., 2008). Signal maps and peak locations were used as input data to Active Motifs proprietary analysis program, which creates Excel tables containing detailed information on sample comparison, peak metrics, peak locations and gene annotations. No normalization was performed on the H3K27me3 data, while standard normalization was applied to the H3K4me3 data. The tag number of all samples (within a comparison group) was reduced by random sampling to the number of tags present in the smallest sample. To compare peak metrics between 2 or more samples, overlapping Intervals were grouped into “Merged Regions”, which are defined by the start coordinate of the most upstream Interval and the end coordinate of the most downstream Interval (= union of overlapping Intervals; “merged peaks”). In locations where only one sample has an Interval, this Interval defines the Merged Region. The sequencing tracks (number of fragments in each 32 bp bin stored as a “bigwig” file) were visualized with the UCSC genome browser.

#### Three germ layer differentiation

The differentiation of bat pluripotent stem cells was carried out with the STEMdiff Trilineage differentiation kit (StemCell Technologies, 05230) following the manufacturer’s protocol. Cells were plated at the desired densities in mTeSR medium (StemCell Technologies, 85850), and plated on Vitronectin-coated (StemCell Technologies, 07180) cell culture plates. After 5 days (endoderm or mesoderm) or 7 days (ectoderm) in culture as directed by the manufacturer. For the ectoderm differentiation, the floating three-dimensional structures were then replated and grown for 4 additional days in fibroblast medium. The cells were stained with antibodies detecting the appropriate lineage markers as described above or cells were collected (surface area of 10 cm^2^ per replicate) for RNA isolation and RNA-seq after addition of 600 µl lysis buffer RTL (part of the RNeasy kit; Qiagen, 74104).

#### Embryonic body differentiation

Bat pluripotent stem cells grown on irradiated mouse embryonic fibroblasts from a total area of 60 cm^2^ were washed with PBS, treated for 10 minutes with Gentle Cell Dissociation Reagent (StemCell Technologies, 07174), collected by centrifugation and resuspended in 12 ml differentiation medium consisting of DMEM/F-12 (Life Technologies, 11330-057), 10% fetal bovine serum (Sigma, F4135), 0.1 mM MEM Non-essential amino acids (Life Technologies, 11140-050), 2 mM GlutaMax supplement (Life Technologies, 35050-061), Penicillin-Streptomycin (10 U/ml and 10 µg/ml, respectively; Life Technologies, 15140122) and 100 µM 2-mercaptoethanol (Fluka, 63689). The cells were then transferred to one uncoated 60 cm^2^ petri dish (Corning, 351029). After 3 days in culture, as much as possible of the medium (about 2/3) was carefully exchanged without disturbing and removing the floating EBs that had formed. The floating EBs were collected after 3 more days (total of 6 days) in culture, fixed in Cytofix/Cytoperm fixation buffer (Becton Dickinson, BDB554714) overnight, and then stained with antibodies against as described above to detect differentiation markers of all three germ-layers by immunofluorescence. For RNA isolation and RNA-seq, EBs were formed as described, collected, resuspended in 6 ml differentiation medium, and distributed into three wells of cell-culture treated 6-well plates (10 cm^2^ each). After 2 more days in culture, the cells were washed with PBS, lysed with 600 µl buffer RTL (part of the RNeasy kit; Qiagen, 74104) and RNA was isolated as described above.

#### Blastoid differentiation

Cells were harvested and plated as described for the embryonic body formation above. After 3 days in culture, 100 ng/ml BMP4 (R&D Systems, 314-BP-010) were added to the medium. 24 later the supernatant was diluted with 2/3 of fresh medium and transferred to two fresh uncoated petri dishes. The medium was exchanged after 3 more days in culture and floating blastoids were harvested 4 days later (total of 12 days of differentiation). The blastoids were fixed in Cytofix/Cytoperm fixation buffer (Becton Dickinson, BDB554714) overnight, and stained as described above to detect the expression of Oct4 by immunofluorescence microscopy.

#### Teratoma formation

Two 6-well plates (12 wells) of bat pluripotent stem cells grown on irradiated mouse embryonic fibroblasts were scraped off in 2 ml DMEM/F-12 medium (Life Technologies, 11330-057), collected by centrifugation and resuspended in 500 µl DMEM/F-12 medium. 100 µl of the cell suspension were injected into the hindleg muscle of 8-week-old male Fox Chase SCID Beige Mice (Charles River, 250). Tumor tissue that had formed after 16 weeks was harvested, fixed in 10% Formalin (Fisher Scientific, SF1004) overnight and then transferred to 70% ethanol. The tissue was submitted to the Pathology Core at the Icahn School of Medicine at Mount Sinai for paraffin embedding and hematoxylin and eosin staining of 5 µm sections. Images were acquired with an AxioObserver microscope (Zeiss).

#### Western Blot analyses

Cells were lysed with RIPA lysis and extraction buffer (Fisher Scientific, 89900) containing Proteinase Inhibitor (Roche, 45582400) for 30 minutes on ice, debris was removed by centrifugation and the supernatant was transferred to a new tube. Protein concentration was determined using the BCA Protein assay kit (Pierce, 23252) following the manufacturers recommendation in 96 well format. 20 µg of protein isolated from each cell line were separated on a 12% TGX Precast gel (Bio-Rad, 4561044) for 70 minutes at 200 V in TGS buffer (Bio-Rad, 1610772), the Precision Plus Kaleidoscope Protein standard (Bio-Rad, 161-3075) was used as size marker. The protein was transferred to a PVDF membrane (activated with methanol) with 1x TGS Buffer (Bio-Rad, 161-0374) supplemented with 20% methanol using the Turbo blot system (Bio-Rad, 1704150) for 30 minutes at 25 V (1 mA). The membrane was blocked with EveryBlot blocking buffer (Bio-Rad, 12010020) for 30 minutes, and then incubated with the anti-Pan Corona (Invitrogen, MA1-82189) or anti-HERVK(cap)(Austral Biologicals, HERM18315) antibodies in a 1:1000 dilution in EveryBlot blocking buffer for 1-hour. The membrane was washed four times with Tris-buffered saline (Fisher Scientific, BP2471500) supplemented with 0.05% Tween 20 (Bio-Rad, 161-0781) for 5 minutes, incubated with the secondary anti-mouse-HRP antibody in a 1:1000 dilution (Promega, PR-W4021) for 1 hour, and then washed as before. Signals were developed with the Clarity Western ECL Substrate (Bio-Rad, 170-5060) and detected with the ChemiDoc MP Imaging System (Bio-Rad, 170-01402).

#### Principal Component Analysis (PCA)

The DESeq2 output files of the RNA-seq analyses described above were subjected to a Variance Stabilizing Transformation (VST) using within-group-variability (Anders and Huber, 2010) to compare the bat pluripotent stem cell transcriptional profile with that of other species. The first two principal components of this result were plotted using the ggscatter function (https://rpkgs.datanovia.com/ggpubr/reference/ggscatter.html) from the R package ggpubr (https://cran.r-project.org/web/packages/ggpubr/index.html), and the weight of each gene’s contribution to the principal components were extracted for PC1 and PC2. The datasets used in the PCA were: GSM4616525, GSM4616526 and GSM4616527 (dog iPS), GSM4617887, GSM4617889, GSM4617890, GSM4617891, GSM4617895, GSM4617900 and GSM4617901 (marmoset iPS), GSM4616532 (human iPS), GSM4616535 and GSM4616536 (pigIPS) from study GSE152493 (Yoshimatsu et al., 2021), and GSM1287734, GSM1287745 and GSM1287746 (mouse ESC) and GSM1287736, GSM1287747 and GSM1287748 (mouse iPS) from GSE53212 (Carter et al., 2014), as well as GSM2718393 and GSM2718399 (mouse iPS) from GSE101905 (Knaupp et al., 2017).

#### Evolutionary selection analysis

To explore evidence of positive selection in *R. ferrumequinum* for the 674 genes identified as part of the “leading” edge in the PCA analysis described above, we extracted all gene alignments that were available for these transcripts (n = 491) and had previously been annotated (Jebb *et al.,* 2020), in addition to annotating 169 alignments that had been made available as part of BAT1K but were currently unannotated. These alignments contained a maximum of 48 species from all eutherian mammalian superorders, with the species tree published by Jebb *et* al. (2020) used for all selection analyses. A total of 660 of these alignments contained representative genes for *R. ferrumequinum* and were analysed for positive selection using the branch-site models in the codeml package of the PAML suite of software (Yang, 2007). Positive selection was inferred using likelihood-derived dN/dS (ω) values under both a null (foreground and background ω constrained to be less than 1) and alternative (foreground ω can vary) model. The *R. ferrumequinum* lineage was designated as foreground branch to detect unique instances of taxon-specific positive selection. A likelihood ratio test (LRT, 2*lnL_alt_–lnL _null_) was used to compare the fit of both models, with a p-value calculated assuming chi-squared distributed LRTs. P-values were corrected for multiple testing using the Benjamin-Hochberg False Discovery Rate (FDR) method via *‘padjust’* implemented in R. Any significant gene showing a p-value greater than 0.05 with ω >1 was explored further. Significant sites showing positive selection were identified using Bayes Empirical Bayes (BEB) scores with a probability > 0.95. All significant genes were subject to a visual inspection of the alignment, to rule out potential false positive results having occurred due to misaligned sequences. In addition to *R. ferrumequinum*, the *Myotis myotis* (n=637 representative genes), *Homo sapiens* (n=652)*, Mus musculus* (n=628), *Canis lupus* (n=593) and *Felis catus* (n=603) lineages were also independently designated as foreground branches for all genes containing a representative sequence shared with *R. ferrumequinum*. This served as a means of determining whether positive selection identified in *R. ferrumequinum* was truly unique to the species lineage or a consequence of bat-specific, Laurasiatherian-specific, or eutherian mammal-specific instances of sequence evolution.

#### Gene ontology and KEGG pathway analyses

Gene ontology and KEGG pathways that are enriched within a group of genes were identified with the Enrichr tool (Xie et al., 2021; maayanlab.cloud/Enrichr/). The odd ratios were then plotted with ggplot2 (Wickham, 2016; cran.r-project.org/web/packages/ggplot2/index.html) with the odds ratio displayed on the x-axis, the dot size reflecting the gene count (number of genes present in the top 5% of PC1 contributing genes) and the dot color reflecting the p-value.

#### Protein Interaction Network

The genes of the Corona virus disease related KEGG pathway were retrieved from the PathCards database (https://pathcards.genecards.org). The differential expression analysis was performed between bat (this study) and mouse iPS cells (GEO accession number: GSM1287736, GSM1287747 and GSM1287748 from Study GSE53212 (Carter et al., 2014) using DESeq2 (Love et al., 2014). The Corona virus disease-related genes were then illustrated with Cytoscape (Version 3.8.2, Shannon et al., 2003) using the STRING protein query with a 0.8 confidence score cutoff. The nodes were colored based on the log2FoldChange with a negative (blue) fold change indicating down-regulation and a positive (red) fold change indicating upregulation in bat pluripotent stem cells cells. Bold borders indicate proteins that were present in the top 5% of PC1 in the PCA analysis described above.

#### Electron microscopy

Cells were grown in chambered Permanox slides (LabTek, 70390) on irradiated mouse embryonic fibroblasts as described above for 5 days and then further processed by the Biorepository and Pathology core at the Icahn School of Medicine at Mount Sinai. Briefly, the cells were rinsed once with DPBS and fixed overnight with 2% paraformaldehyde and 2% glutaraldehyde in 0.01 M sodium cacodylate buffer at 4°C. Sections were rinsed in 0.1 M sodium cacodylate buffer, followed by a quick rinse with ddH_2_O. Cells were post fixed with 1% aqueous osmium tetroxide for 1 hour, followed with an En bloc stain of 2% aqueous uranyl acetate for 1 hour. Sections were washed again in ddH_2_O, dehydrated through graduated ethanol (25-100%), infiltrated through an ascending ethanol/epoxy resin mixture (Embed 812, EMS), and then covered with pure resin overnight. Chambers were separated from the slides, and a modified #3 BEEM embedding capsule (EMS) was placed over defined areas containing cells. Capsules were filled with pure resin and placed in vacuum oven to polymerize at 60 °C for 72 hours. Immediately after polymerization, the capsules were snapped from the substrate to dislodge the cells from the slide. Semithin sections (0.5 – 1 µm) were obtained using a Leica UC7 ultramicrotome (Leica, Buffalo Grove, IL), counterstained with 1% Toluidine Blue, cover slipped and viewed under a light microscope to identify successful dislodging of cells. Ultra-thin sections (85 nms) were collected on 300 hexagonal mesh copper grids (EMS) using a Coat-Quick adhesive pen (EMS). Sections were counter-stained with uranyl acetate and lead citrate and imaged with a Hitachi 7700 Electron Microscope (Hitachi High-Technologies) using an advantage CCD camera (Advanced Microscopy Techniques). Images were adjusted for brightness, contrast, and size using Adobe Photoshop CS4 11.0.1.

#### Image-based flow cytometry (ImageStream)

Cells were seeded onto 6-well plates and separated from irradiated MEFs via two-stage trypsinization after four days. Wells were dosed and incubated with 0.25ml prewarmed (37°C) trypsin which was removed and discarded at 4 minutes. An additional 0.25ml trypsin was added and the plate was again incubated. After eight minutes cells were removed and pelleted via centrifugation. The cells were washed twice in PBS containing 0.5% BSA (Sigma), fixed and permeabilized with Cytofix/Cytoperm (Invitrogen, C10337). The Primary antibody was added at a dilution of 1:200 in wash buffer incubated overnight at 4°C. The cells were washed twice with 0.5% BSA/PBS, resuspended in wash buffer containing the secondary antibody at a 1:200 dilution Cells were then resuspended in wash buffer, the secondary goat anti-mouse AF568 antibody (Life Technologies) and incubated for 1 hour at 4°C. The cells were washed as before resuspended in 0.5% BSA/PBS containing two drops/ml DyeCycle Violet to stain the nuclei.

Imaging was conducted with the ImageStream MkII, at 60x magnification with the extended depth of field mode for probe resolution. Images were acquired using the INSPIRE 2.0 software at the lowest flow speed. Fluorophores were excited by the 405 nm and 568 nm lasers at 60mW and 100 mW, respectively. Cells in focus were gated via histogram of brightfield gradient R.M.S. values and an aspect ratio vs. area plot was used to select the population of single cells. 5000 individual images of focused single cells were taken. Gating was refined further post-acquisition via the IDEAS 6.2 software suite by the same methods and plots, yielding n=1846 (BiPS). This software was used also for image processing, in which a set of custom masks defined by logical operators were used to denote vesicles and sensitively assess probes. For vesicles, it was observed that they may be selected from other cell component by contrast (bright and dark) and also by aspect ratio, and therefore are defined here by “Dilate(Range(Dilate(Range(System(Peak. (Threshold(M01, BF, 70), BF, Bright, 1), BF, 20), 0-5000, 0.4-1), 1), 0-5000, 0.4-1), 1) Or Range (AdaptiveErode(LevelSet(M01, BF, Dim, 5), BF, 75), 0-5000, 0.5-1).” BF and BF2 represent each brightfield image taken of a single cell from each of the two cameras, M01 and M09 represent the corresponding channel masks for each channel and the remaining terms represent mask modifiers and their associated values in the IDEAS software. For resolving immunofluorescence, “Peak(System(M05, Ch05, 3), Ch05, Bright, 1)” where Ch05 represents the staining of interest and M05 represents the corresponding channel mask. Modification was necessary to sensitively include all representative fluorescence, and to distinguish individual foci. The nuclear mask corresponding to DyeCycle Violet staining was defined “Object(M07, Ch07, Tight)” and the cytoplasm was defined through subtraction of the nuclear and vesicle masks from the cell mask through the logical operator available in the software (“Not”). Vesicle-nucleus overlap was determined in favor of vesicles by excluding them from the nuclear mask (“Not”). Probe localization was then defined according to these entities using the respective definitions and the operator “And.” Statistics for foci were generated using the Spot Count feature with a connectedness of 4. Prism 9 was used for graphs and statistics.

#### Retrovirus assay

2 ml of tissue culture medium were collected, and retroviral particle concentrations were determined using the QuickTiter Retrovirus Quantitation Kit (Cell Biolabs, VPK-120) according to the manufacturer’s instructions.

#### Reverse Transcriptase Assay

Reverse transcriptase enzyme levels were determined with the colorimetric reverse transcriptase kit (Roche) per the manufacturer protocol. Cells lines represented were lysed in RIPA buffer, frozen at -80°C, thawed on ice, collected and resuspended in the kit lysis buffer (10 µL pellet in 40 µL lysis buffer per colorimetric well). Incubation duration (15h at 37°C) was selected for maximal sensitivity to the limit of the kit (1-5 pg RT). Absorbance at 405nm was measured by microtiter ELISA plate reader. Sample absorbance measurements were fitted to a linear regression of the measured HIV-1 RT standards (Y=2.549X) to obtain RT concentrations in units of ng/well.

#### Plaque Assay

Supernatants were centrifuged at 10000 rpm for 5 min to remove cellular debris, and the cleared lysates transferred to new tubes. Lysates were then diluted in 10-fold dilutions 6 times. Quantification of infectious titer was then performed by plaque assays in comparison to SARS-CoV-2 infection as positive control. Briefly, Vero-E6 cells were plated as confluent monolayers in 12 well dishes. Media was removed, and wells washed in 1ml of PBS. 200ul of diluted lysates was then added per well and allowed to incubate for 1 hour at 37°C. After viral adsorption, lysates were removed from the well and cells were overlaid with Minimum Essential Media supplemented with 2% FBS, 4 mM L-glutamine, 0.2% BSA, 10 mM HEPES and 0.12% NaHCO3 and 0.7% agar. 72h post infection, agar plugs were fixed in 10% formalin for 24h before being removed. Plaques were visualized by staining with TrueBlue substrate (KPL-Seracare) and viral titers calculated and expressed as PFU/ml.

#### Metapneumovirus (MPV) infection of BiPS and mES cells

50,000 mouse ES cells (R1) or BiPS cells were plated per well of a 12-well plate on irradiated CF1 mouse embryonic fibroblasts using mouse and bat culture medium respectively. After 24 hours, culture medium containing human Metapneumovirus with GFP (MPV-GFP) (ViralTree, M121) with a final multiplicity of infection (MOI) of 3. Medium was changed daily, and samples were dissociated at 3 and 5dpi using trypsin/EDTA and the infection rate was determined by fluorescence activated cell sorting (FACS)

#### Iso-Seq library preparation and sequencing

Cells at passage 27 were lyzed in 400 µl Trizol reagent (Life Technologies, 15596-018) and total RNA was extracted using the AllPrep DNA/RNA Mini Kit (Qiagen, #80204) including a DNase digest to remove any potential contamination from carryover of genomic DNA using RNase-free DNase (Qiagen, 79254) according to the manufacturer’s instructions. The extracted RNA was then purified using 1.8X RNAClean XP beads (Beckman Coulter) to remove any molecular impurities. Iso-Seq SMRTbell libraries were prepared as recommended by the manufacturer (Pacific Biosciences). Briefly, 300 nanograms of total RNA (RIN > 8) from each sample was used as input for cDNA synthesis using the NEBNext Single Cell/Low Input cDNA Synthesis & Amplification Module (NEB, E6421L), which employs a modified oligodT primer and template switching technology to reverse-transcribe full-length polyadenylated transcripts. Following double-stranded cDNA amplification and purification, the full-length cDNA was used as input into SMRTbell library preparation, using SMRTbell Express Template Preparation Kit v2.0. Briefly, a minimum of 100 ng of cDNA from each sample were treated with a DNA Damage Repair enzyme mix to repair nicked DNA, followed by an End Repair and A-tailing reaction to repair blunt ends and polyadenylate each template. Next, overhang SMRTbell adapters were ligated onto each template and purified using 0.6X AMPure PB beads to remove small fragments and excess reagents (Pacific Biosciences). The completed SMRTbell libraries were further treated with the SMRTbell Enzyme Clean Up Kit to remove unligated templates. The final libraries were then annealed to sequencing primer v4 and bound to sequencing polymerase 3.0 before being sequenced on one SMRTcell 8M on the Sequel II system with a 24-hour movie each. After data collection, the raw sequencing subreads were imported to the SMRTLink analysis suite, version 10.1 for processing. Intramolecular error correcting was performed using the circular consensus sequencing (CCS) algorithm to produce highly accurate (>Q10) CCS reads, each requiring a minimum of 3 polymerase passes. The polished CCS reads were then passed to the *lima* tool to remove Iso-Seq and template-switching oligo sequences and orient the isoforms into the correct 5’ to 3’ direction. The *refine* tool was then used to remove polyA tails and concatemers from the full-length reads to generate final full-length, non-chimeric (FLNC) isoforms. The FLNC isoforms were then clustered together using the *cluster* tool to generate final, polished consensus isoforms per sample.

#### Identification and illustration of viral sequences in the bat pluripotent stem cell transcriptome

The existence of viruses in the *Rhinolophus ferrumequinum* transcriptome was explored by analyzing the RNA-seq and Iso-seq data based on a metagenomic approach using Kraken2 v2.1.2 (Wood et al, 2019). First, the adaptors in the RNA-seq data were removed with Trimgalore v0.6.7 (Krueger et al., 2021) and all replicates for corresponding datasets were joined in one file. The reference library “RefSeq complete viral genomes / proteins” was downloaded and a custom database was built to identify matches within the processed RNA-seq or Iso-seq. To eliminate false positive hits that could be due to matches with any cellular transcript such as oncogenes that are carried by some viruses, a second analysis was performed after eliminating all reads from the RNA-seq and Iso-seq datasets that matched any annotated *Rhinolophus ferrumequinum* transcript. To do this, the Iso-Seq FLNC isoforms or RNA-seq trimmed fastq sequences were first mapped to the “*Rhinolophus ferrumequinum* genomic rna exons RefSeq” file “GCF_004115265.1_ mRhiFer1 _v1.p_rna_from_genomic.fna” using gmap/gsnap (https://doi.org/10.1093/bioinformatics/bti310). The sequences with no mappings were then used to identify viral sequences using Kraken2 as before.

In a first approximation, we mapped the sequencing reads against a virus database, using a metagenomic classification tool (Kraken2). The results revealed a taxonomically highly diverse “zoo” of assigned viruses belonging to several major viral families. We also classified viral sequences using RNA-seq data from BEFs (Tables S4C and S4D) and from human ES cells (Tables S4E and S4F) notably yielding some viral sequences albeit to a lesser degree (Figure S11B). This observation in BEFs was surprising as post-implantation tissues typically do not exhibit endogenous viral activity^49^, underscoring the notion that bat cells harbor pro-viral environments. Further, the results support our hypothesis that the epigenetic reprogramming of bat fibroblasts reactivates a considerable number of additional dormant viral sequences.

Further, we investigated potential confounding effects that might impact interpretation of the metagenomic data. Three potential sources for distortion are (*i*) statistical stringency, (*ii*) cellular genes containing viral-like sequences (e.g., oncogenes), and (*iii*) potential *xeno* sequence pollution originating from the feeder cells. To address the first point, we utilized progressively higher statistical stringency, yielding an expected decrease in matches in our *R. ferrumequinum* RNA-seq data. However, even with the most stringent parameters tested, there remained a sizable number of hits (Table S4E). To exclude potential cellular genes misinterpreted by the classification algorithm as viruses, we depleted the RNA-seq and Iso-seq from all sequences that match exons, which only marginally affected the number of hits (Table S4I, S4J, and S4K). When the Kraken2 database was expanded to include bacteria, archaea, and the whole human genome, the yield was only 0.65 percent, compared to 2.53 percent when the viral database was probed (Figure S11D). Finally, we checked if some of the classified sequences were of murine origin and found that this was the case for several retroviruses, and for example, the sequences classified as Diatom colony associated ssRNA virus 1. The latter matched a genomic region on mouse chromosome chr9:65,969,137-65,969,198 (GRCmm39/mm39).

#### Mapping of RNA-seq reads to bat genomes and quantifying expression of ERVs

To trim adapters and generate quality metrics of the fastq files, we used Trimmgalore v.0.6.6 (https://github.com/FelixKrueger/TrimGalore), a wrapper for Cutadapt (https://github.com/marcelm/cutadapt) and FastQC (https://www.bioinformatics.babraham.ac.uk/projects/fastqc/). Then, reads were mapped to the genome of *R. ferrumequinum* (Bat1K assembly HLrhiFer5) using HISAT2 v.2.2.1 (Kim et al. 2019) suppressing unpaired alignments for paired reads (--no-mixed), suppressing discordant alignments for paired reads (--no-discordant), and setting a function for the maximum number of ambiguous characters per read (--n-ceil L,0,0.05). Output files were then filtered to remove any unmapped reads (-F 4), sorted and indexed using samtools (Li et al., 2009). Aligned reads were then assembled into transcripts using stringTie v2.2.1 (Pertea et al. 2015) in stranded mode (-rf). To generate a Ballgown readable expression output with normalized expression units of fragments per kilobase of transcript per million mapped fragments (FPKMs), we also used as input in strigTie the Bat1K annotation of known endogenous retrovirus (ERVs) for *R. ferrumequinum* (Jebb et al. 2020) (https://genome.senckenberg.de/). Output counts were post-process and plotted with a custom R script.

#### *De novo* assembly of potential virus-derived RNA-seq

The trimmed reads that were identified by Kraken2 v2.1.2 to map to viral sequences with a confidence score of 0 as described above were first grouped together based on their taxonomic ID assigned by Kraken and then classified as either mammalian or non-mammalian using the VIRION database (Carlson et al., 2022). The data were converted to FASTA format using the Seqtk v1.3 program and the reads were assembled using the Trinity v2.12 software. To check and gather successful assemblies that had produced at least one contig, a custom BASH script was applied for both groups of mammalian and non-mammalian viruses.

#### Mapping transcripts to viral and mammalian databases

To determine if the assembled transcripts represented an expressed viral sequence, all transcripts were mapped to a database of viral genomes using BLAST. The viral database consisted of genomes whose host species contained either ‘human’ or ‘vertebrate’ as specified in the NCBI database. Initially this list contained over 17,000 genomes. However, this was reduced to 3,922 genomes by taking only unique virus/strain names. An additional non-mammalian virus database was generated by combining all genomic sequences of viruses identified by Kraken2 and classified as non-mammalian via VIRION. Transcripts were also mapped to a combined database of bat, human and mouse genomes to both confirm their presence in the bat and to exclude the possibility of false positives through contamination. For each of these transcripts, expected values for both bat and viral genome BLAST results were combined into a single metric via the following formula: Log (bat-expected value+1 x virus-expected value+1). A threshold of less than 0.3, representing a combined e-value of less than 1e^-50^ for both viral and bat hits, was used to rule out potential false positives. In addition, we used SQUID (http://eddylab.org/software.html) to shuffle the 63 (bottom-up) and 82 (top-down) sequences while preserving the dinucleotide distribution (parameter -d) to obtain a conservative threshold to distinguish bona fide viral homology from matches by random chance. Shuffled sequences were mapped to both the bat genome and viral genome databases, with the same BLAST threshold applied. All transcripts passing this threshold were extended by 5000 bp flanks within the bat genome and these regions were subsequently mapped to the viral database to confirm their presence in a viral genome.

---

**Figure S1.**
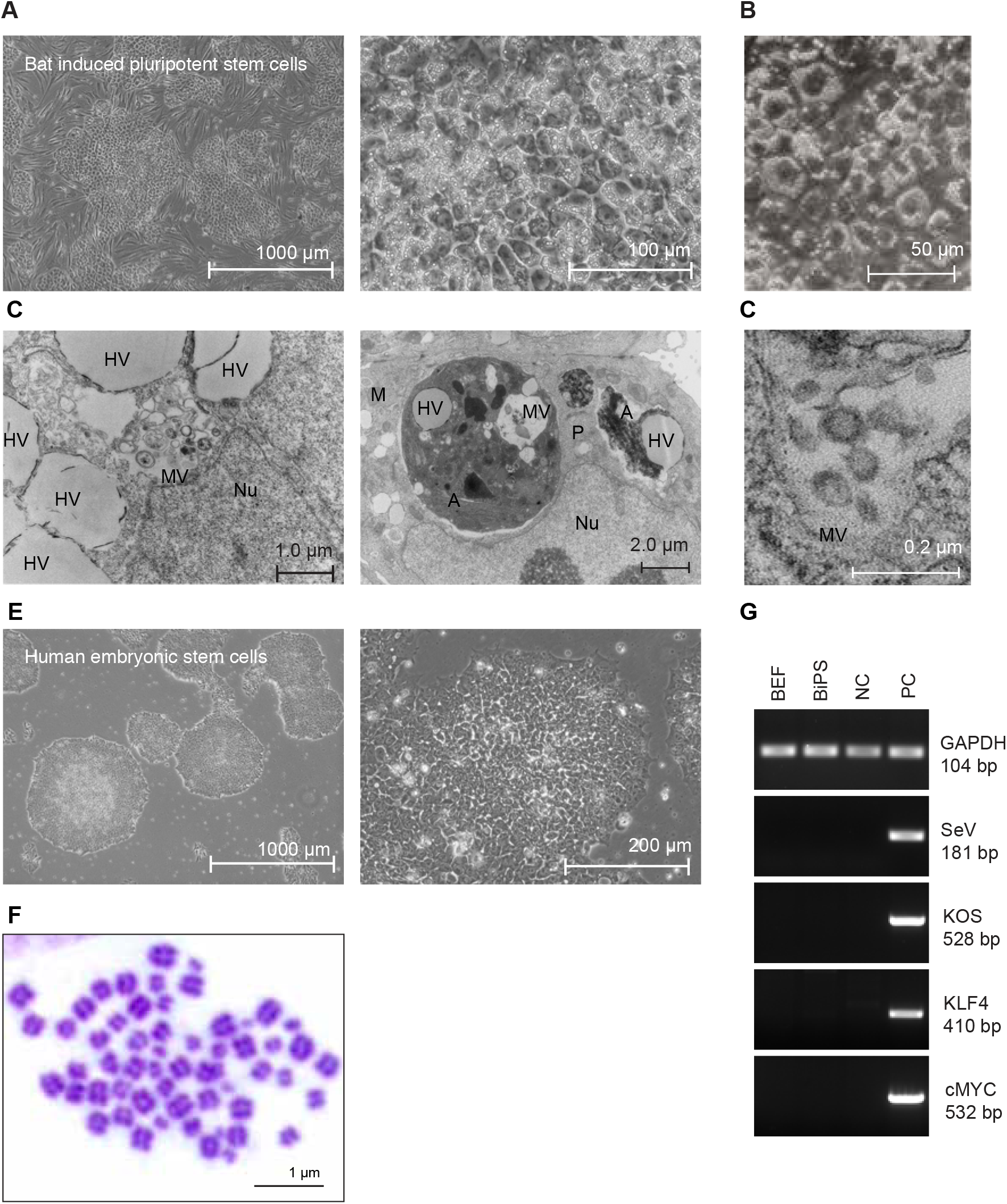
Characterization of pluripotent stem cells generated from *Rhinolophus ferrumequinum* fibroblasts. **(A)** Microscopic images of bat pluripotent stem cells at different magnifications. **(B)** Differential interference contrast microscopy image of BiPS cells highlighting prominent cytoplasmic vesicles. **(C)** Overview of transmission electron microscopy of bat pluripotent stem cells. MV: vesicles filled with multimembrane structures; HV, other vesicle structures filled with homogenous content; Nu, Nucleus; A, autophagosome; M, mitochondria. **(D)** Higher magnification of electron microscopy images as in (c) showing the presence of aggregates that are morphologically compatible with the appearance of endogenous retrovirus-like particles (VLP). **(E)** Microscopic images of human embryonic stem cells (H9) and bat pluripotent stem cells at different magnifications. **(F)** Karyotype analysis of BiPS cells at passage 17. Shown is a representative image after Giemsa staining of metaphase spreads with 56 chromosomes. **(G)** PCR verification of reprogramming-associated virus clearing. Bat iPS cells (BiPS) at passage 92 were tested for Sendai virus clearance in comparison to the embryonic fibroblasts used as starting material (BEF), adult fibroblasts as negative control (NC), and freshly transduced cells at passage 3 as a positive control (PC). bp, base pairs; SeV, Sendai virus; KOS, KLF4-OCT4-SOX2.

**Figure S2.**
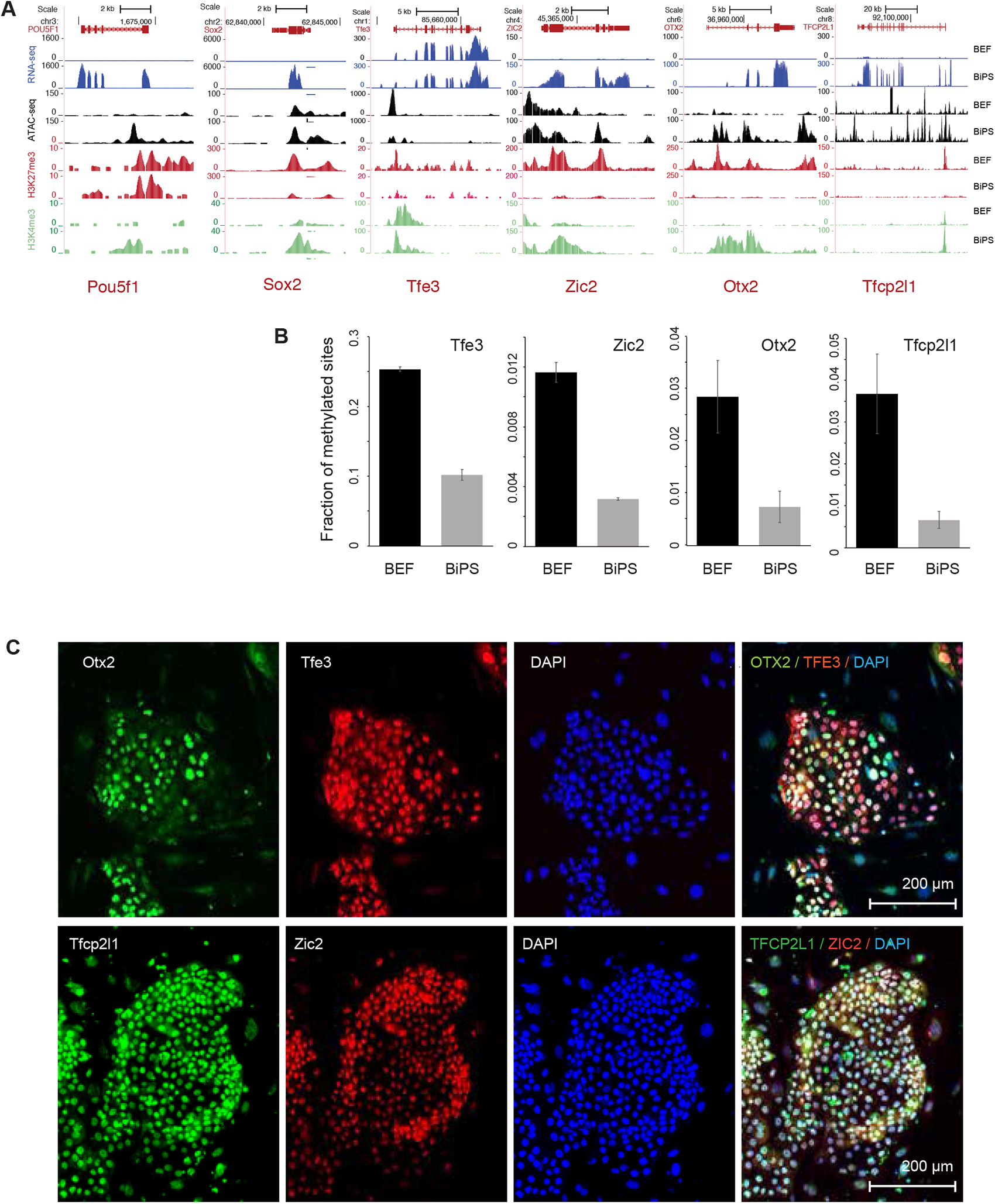
Characteristics of pluripotency markers in pluripotent stem cells generated from *Rhinolophus ferrumequinum* fibroblasts. **(A)** Sequencing tracks showing expression, ATAC-seq signal, Histone H3K27 trimethylation (H3K27me3) and Histone H3K4 trimethylation (H3K4me3) status of pluripotency markers Oct4, and Sox2 in bat embryonic fibroblasts (BEF) or induced pluripotent stem cells (BiPS). **(B)** Fraction of methylated sites in promoters of pluripotency genes that did show promoter methylation. Note that we did not detect methylation in the promoters of Nanog, Pou5f1, or Sox2, which might be related to under-annotation of the *Rhinolophus ferrumequinum* genome at this point in time. **(C)** Immunofluorescence images of bat pluripotent stem cells after staining of markers of naive (Tfe3 and Tfcp2l1) or primed pluripotency (Zic2 and Otx2).

**Figure S3.**
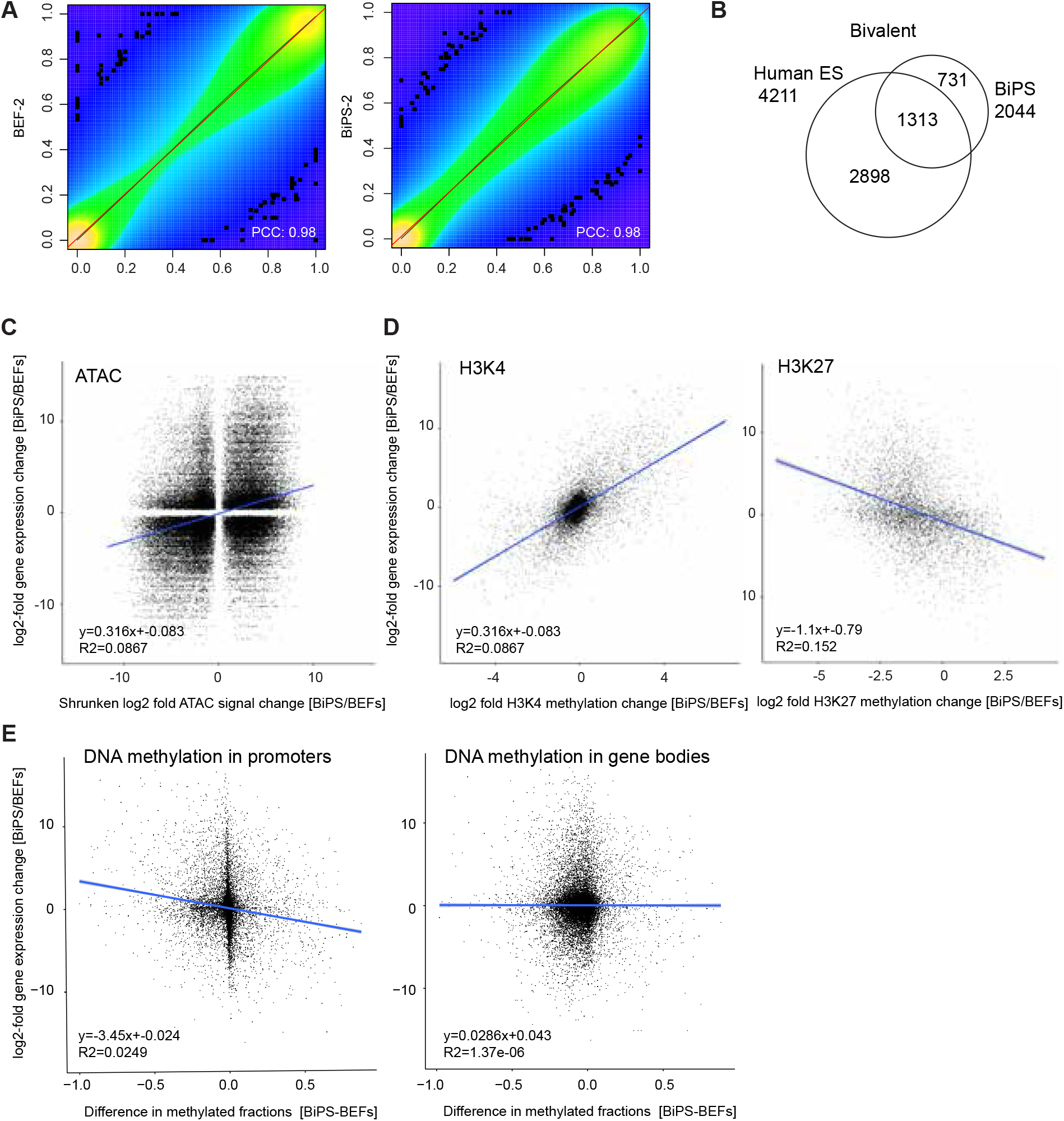
Correlation between epigenetic and transcriptional changes. **(A)** Correlation scatter plot of methylation level at common CpG sites in duplicate samples of BEF or BiPS cells. BEF, bat embryonic fibroblast cells; BiPS, bat pluripotent stem cells; PCC, Pearson correlation coefficient. **(B)** Venn diagram illustrating the overlap of bivalent genes in bat iPSCs and human ES cells. **(C)** Correlation plot of shrunken log2-fold changes in ATAC-seq signal with log2-fold expression changes. Shown are all values with p<0.05. **(D)** Correlation of log2-fold changes in H3K4 trimethylation (H3K4me3, left) or H3K27 trimethylation (H3K27me3, right) with log2-fold changes in gene expression. **(E)** Correlation of log2-fold gene expression changes with the difference in the methylated fraction of promoters (left) or gene bodies (right) fractions.

**Figure S4.**
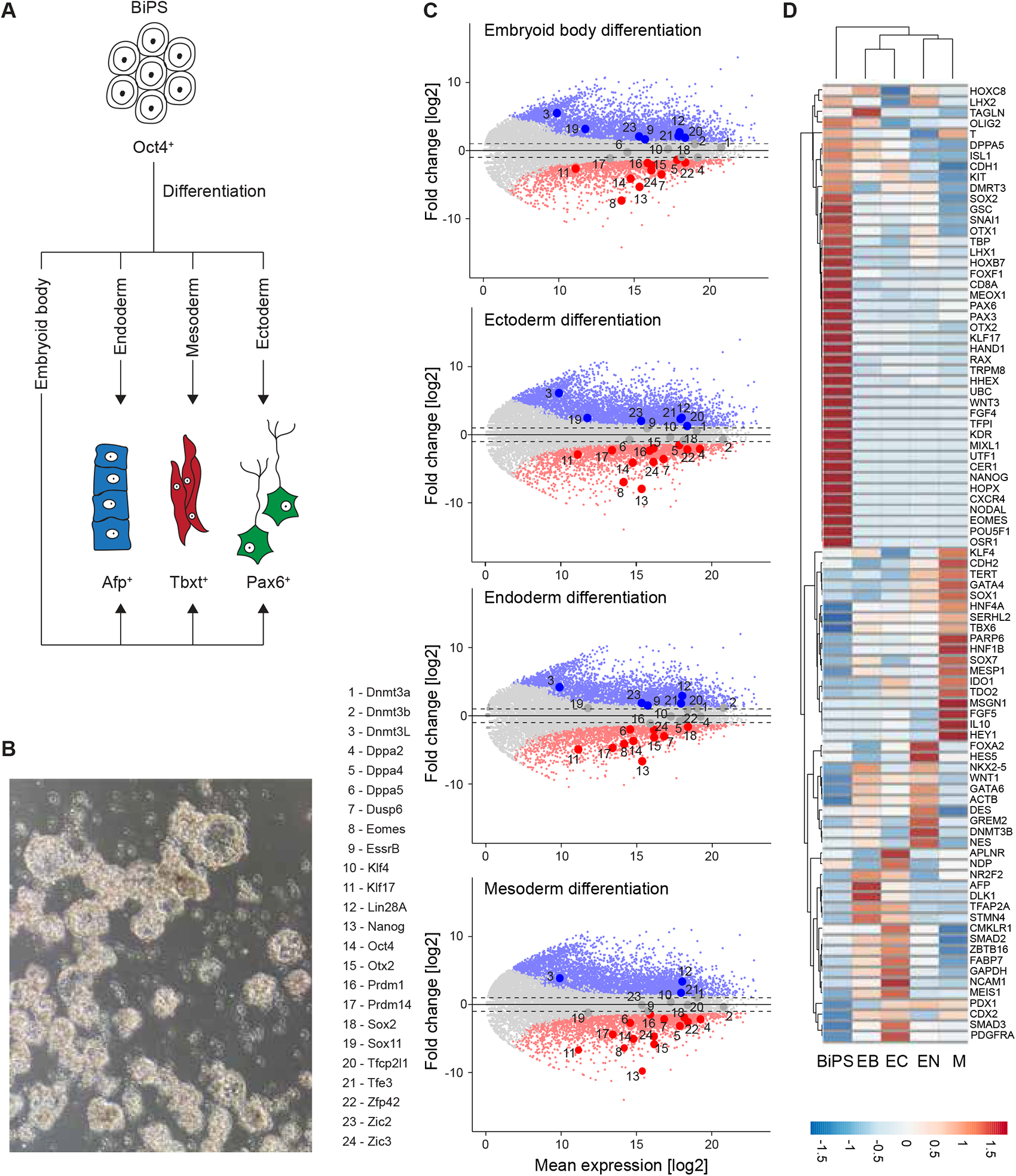
Differentiation potential of bat pluripotent stem cells. **(A)** Schematic of differentiation strategies. **(B)** Phase contrast image of BiPS2-derived embryoid bodies after 7 days of differentiation. **(C)** MA plots depicting the log2 mean expression and log2-fold expression changes of all genes in bat induced pluripotent stem cells (BiPS) after exposure to the noted differentiation conditions illustrated in (A). EB, Embryoid body differentiation; EC, human ectoderm differentiation conditions; EN, human endoderm differentiation conditions; M, human mesoderm differentiation conditions. **(D)** Heatmap depicting expression changes of genes that are known as markers for human ectoderm, mesoderm, or endoderm during the differentiation of BiPS under the conditions described in (A).

**Figure S5.**
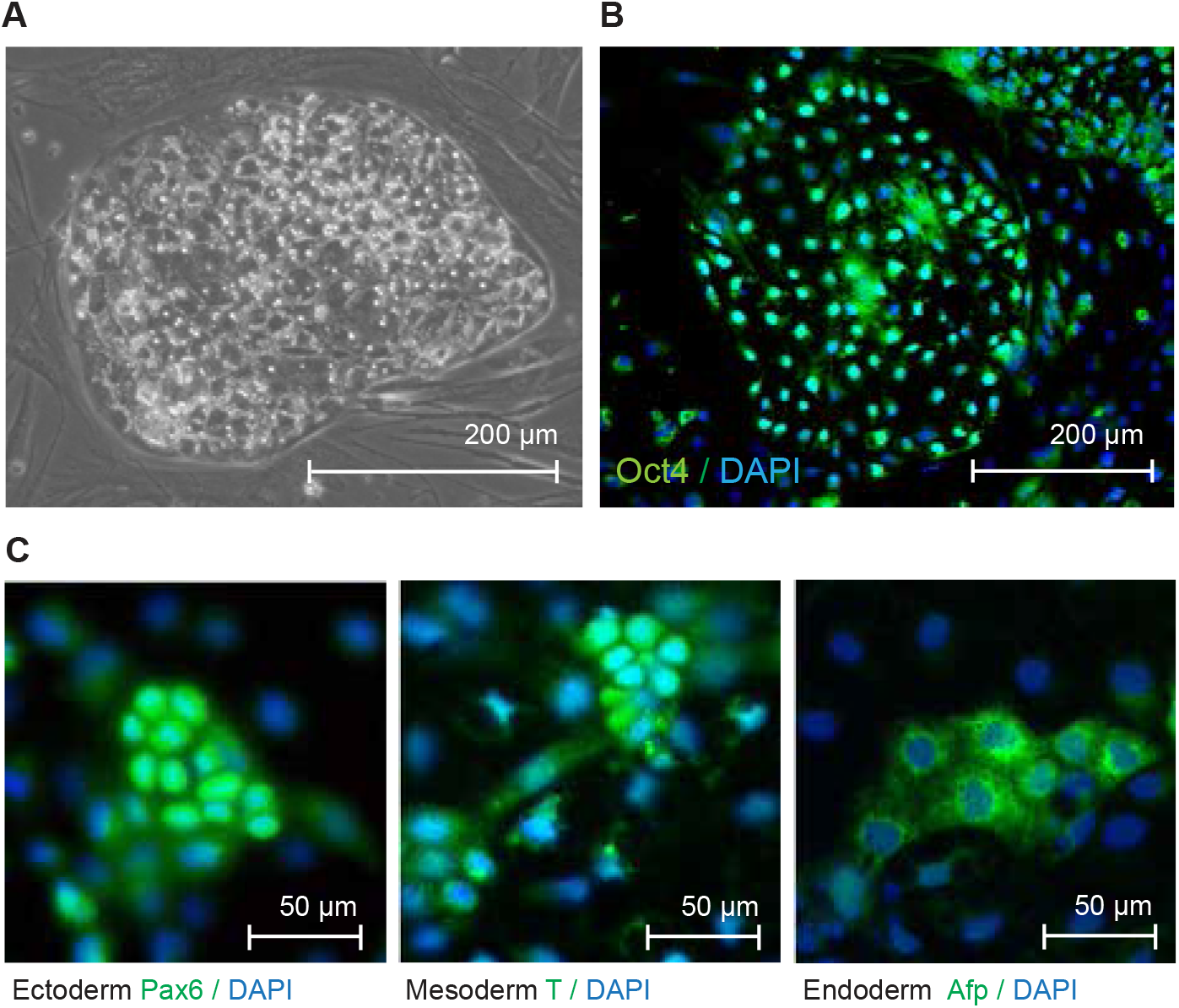
Characterization of *Myotis myotis* induced pluripotent stem cells. **(A)** Phase contrast image of *Myotis myotis* iPS cells. **(B)** Microscopic image of *Myotis myotis* iPS cells after immunostaining to detect pluripotency marker Oct4. **(C)** Microscopic images of *Myotis myotis* iPS cells that underwent differentiation and immunostaining to detect Pax6, Brachyury (T) and Afp as markers for ectoderm, mesoderm and endodem, respectively.

**Figure S6.**
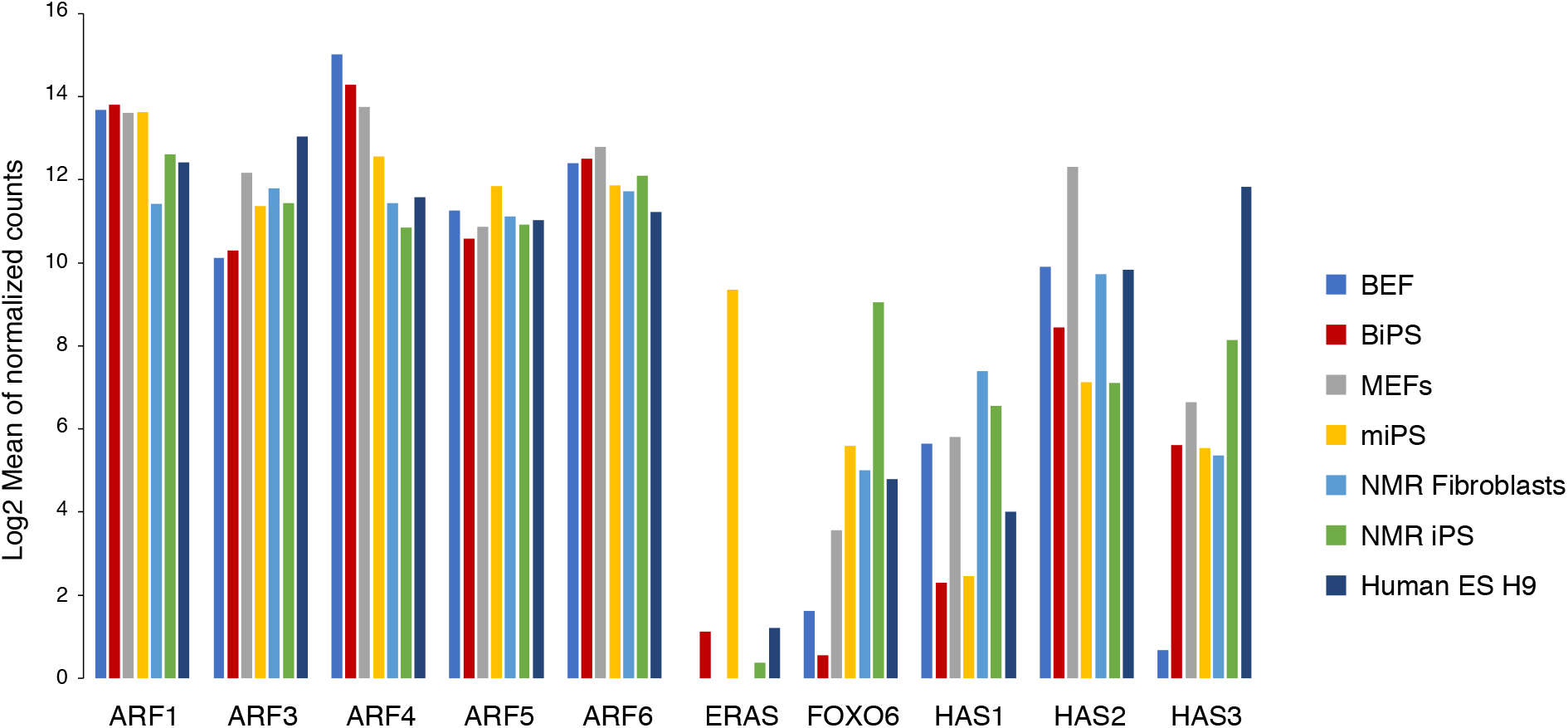
Expression profile of genes associated with tumor suppression. The data sets were from this study (bat), GSE53212 (mouse, GEO), PRJNA400257 (Naked mole-rat, BioProject), and GSE175070 (human, GEO). ARF, ADP ribosylation factor; BEF, bat embryonic fibroblasts; BiPS, bat induced pluripotent stem cells, ERAS, ES cell-expressed Ras; FOXO6, Forkhead Box O6; H9, human ES cells; HAS, Hyaluronan synthase; MEFs, mouse embryonic fibroblasts; NMR, naked mole-rat.

**Figure S7.**
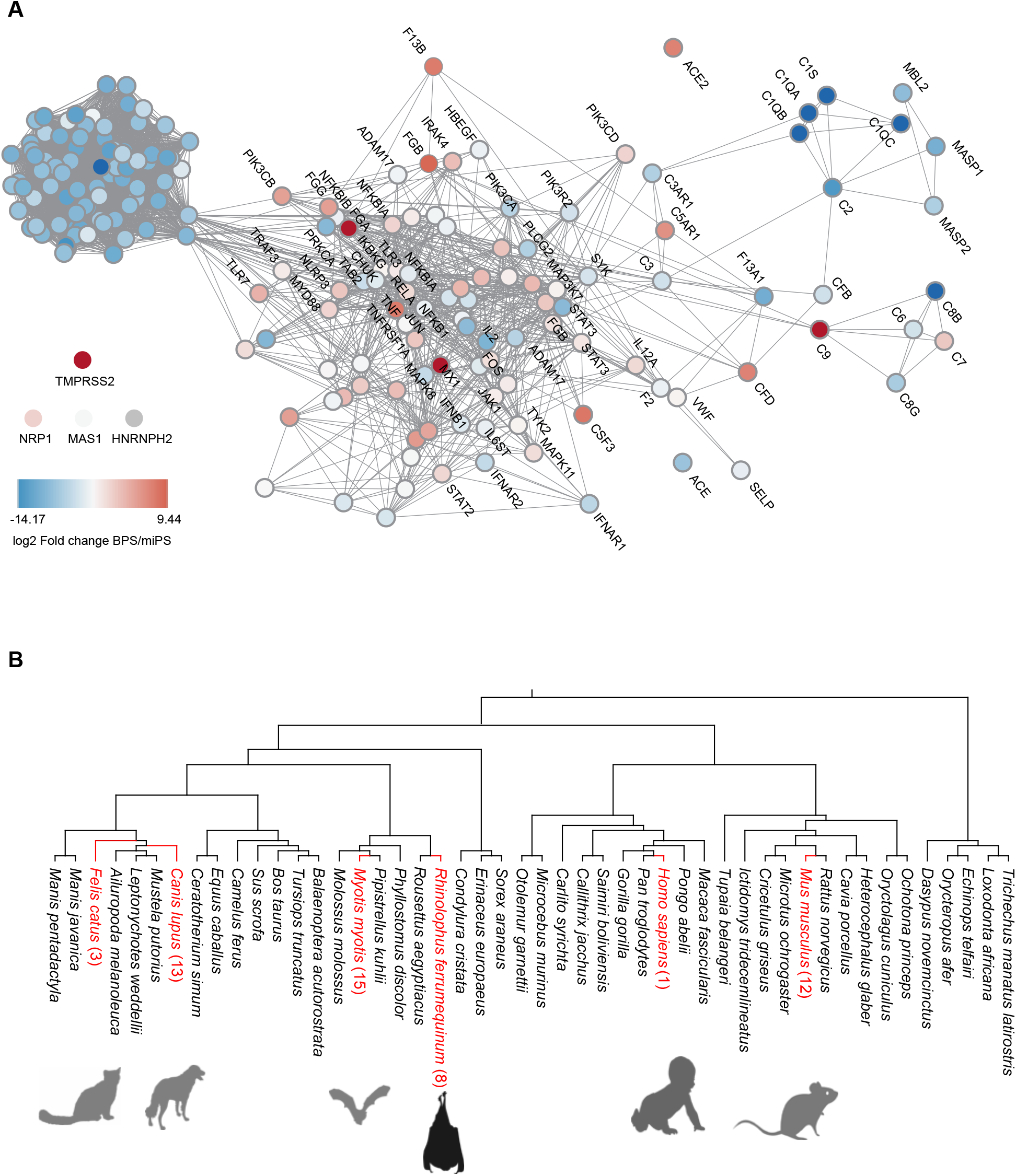
Comparative transcriptomics and evolutionary selection analyses. **(A)** Interaction of genes that are part of the KEGG Corona Virus Disease pathway. Nodes are colored based on the log2-fold change between bat and mouse iPS cells. Red indicates genes expressed at a higher level in BiPS, and blue indicates those expressed at a lower level. Bold borders indicate proteins present in the top 5% of genes in PC1 (leading edge). **(B)** Selection analyses of leading edge-genes by comparative genomics of the *R. ferrumequinum* lineage identified only eight genes (AARD, COL3A1, FAM111A, LAMB3, MUC1*, NES*, RGS5, RSPH1*) with significant evidence of positive selection, five of which showed at least one highly probable BEB site with no visual issues in the alignment region, while three genes (designated with ‘*’) did not. Additional lineages and the number of leading edge-genes with significant evidence of positive selection found in them are highlighted in red.

**Figure S8.**
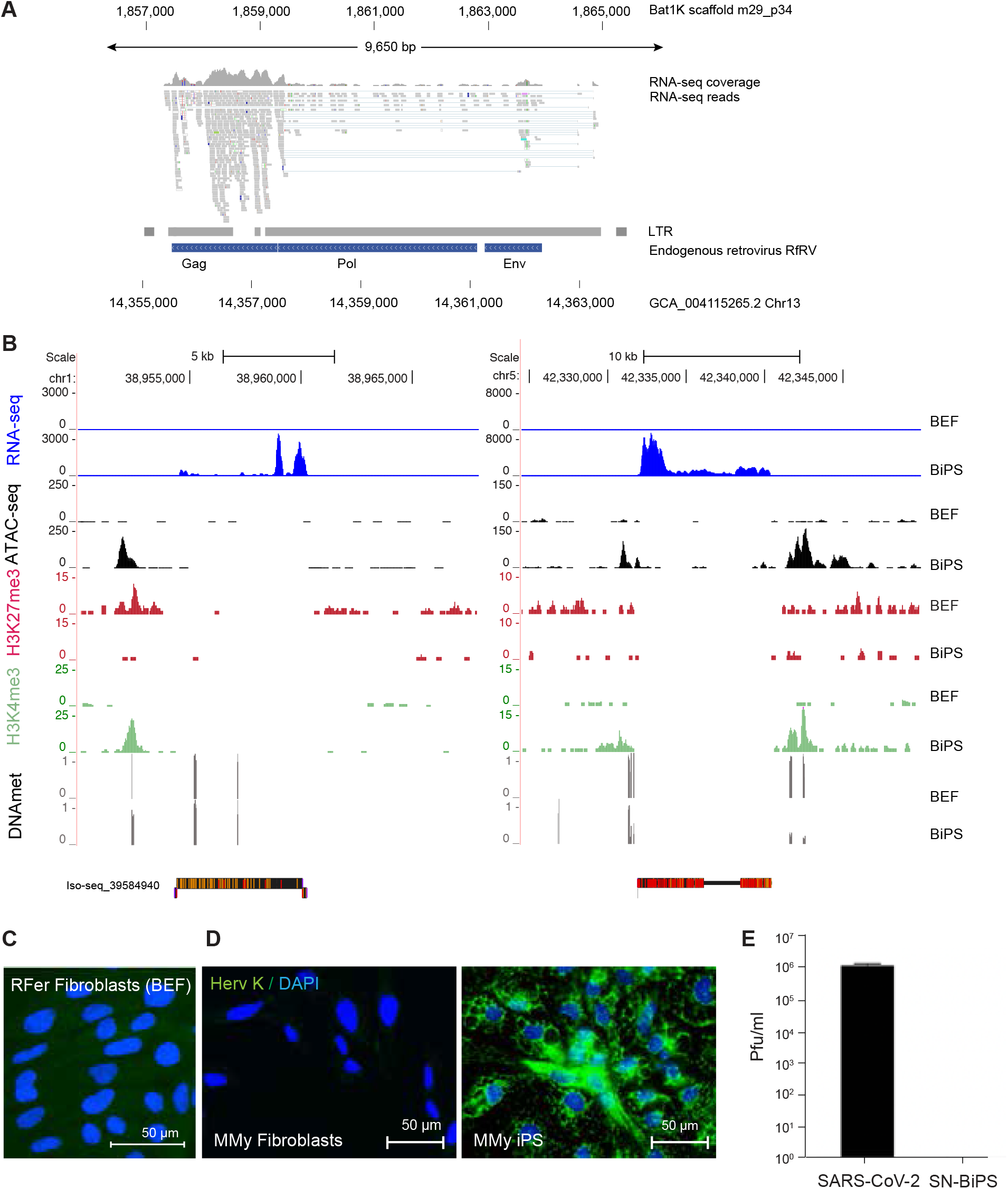
Viral activity in BiPS cells. **(A)** RNA-seq tracks aligning with the genomic *Rhinolophus ferrumequinum* region (Bat1K annotated scaffold, top, and GCA assembly, bottom) containing the previously identified endogenous retrovirus RfRV. LTR, long terminal repeat. **(B)** Sequencing tracks showing expression, ATAC-seq signal, Histone H3K27 trimethylation (H3K27me3), Histone H3K4 trimethylation (H3K4me3) as well as DNA methylation (DNAmet) status in the genomic region on chromosome 1 surrounding RFe-V-MD1 shown in Figure 4 (left) and a genomic region on chromosome 5 that contains a sequence highly similar to RFe-V-MD1 (right). Note that the 6088 bp-long Iso-seq fragment aligns with the (-) strand on chromosome 1 and the (+) strand on chromosome 5. **(C)** Immunostaining with an antibody detecting the endogenous retrovirus protein Herv K (left) in *Rhinolophus ferrumequinum* embryonic fibroblast (BEF) cells. **(D)** Immunostaining as in (C) using *Myotis myotis* fibroblasts (left) or induced pluripotent stem cells. **(E)** Plaque assay with supernatant of *Rhinolophus ferrumequinum* induced pluripotent stem cells compared to active SARS-CoV-2 viral particles in Vero cells. BEF, Bat embryonic fibroblasts; MMy, *Myotis myotis;* RFer, *Rhinlophus ferrumequinum*.

**Figure S9.**
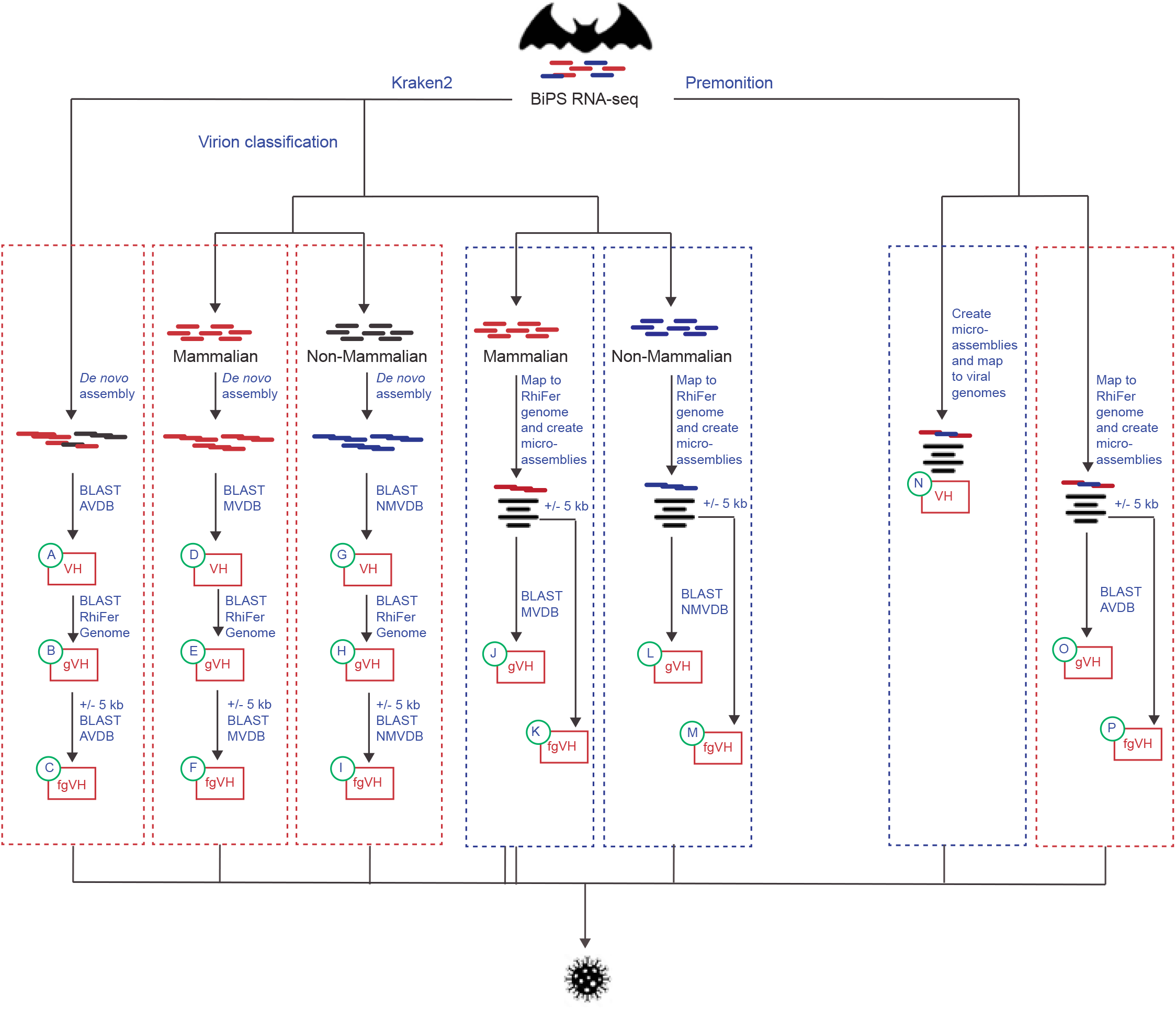
Virome mining approaches. RNA-seq reads were first classified using Kraken2 (*left*). The classified reads were then either analyzed directly or separated by their homology to a mammalian or a non-mammalian virus using VIRION. In a “bottom-up” approach (*red boxes*), the reads that were classified to belong to the same virus were assembled *de novo* and blasted against databases containing all viruses (AVDB), mammalian viruses (MVDB) or non-mammalian viruses (NMVDB). The identified sequences with viral hits were then further mapped to the *Rhinolophus ferrumequinum* (RhiFer) genome (gVH) and in a final step, reads/assemblies that did map to the genome were extended by 5 kb on both sides and the sequences again blasted against the databases as before to extend the search for viral hits in flanking genomic regions (fgVH). In a parallel “top-down” approach (*blue boxes*), the Kraken2 reads were mapped directly to the *Rhinolophus ferrumequinum* genome, microassemblies were generated based on the mapped reads, and blasted directly against the different databases as before or first extended by 5 kb on both sides. In an orthogonal approach, the reads were similarly assigned to viruses using the Microsoft Premonition software (*right*). First all reads were classified as aligning to a virus or as being unaligned, micro-assemblies were generated and mapped to viral genomes (*red box*). Consensus sequences were extracted and blasted against the *R. ferrumequinum* genome, extended by 5 kb and blasted against a database containing all viruses as before. In another “top-down” approach (*blue box*), the Premonition classified reads were mapped to the *R. ferrumequinum* genome first, and microassemblies generated that were blasted against the “All virus” database either directly or after being extended by 5 kb on both sides. AVDB, all viruses database; MVDB, mammalian virus database; NMVDB, non-mammalian virus database; VH, viral hits; gVH, viral hits with homology to the bat genome; fgVH, viral hits using flanked genomic regions; RhiFer, *Rhinolophus ferrumequinum*; kb, kilobase pairs.

**Figure S10.**
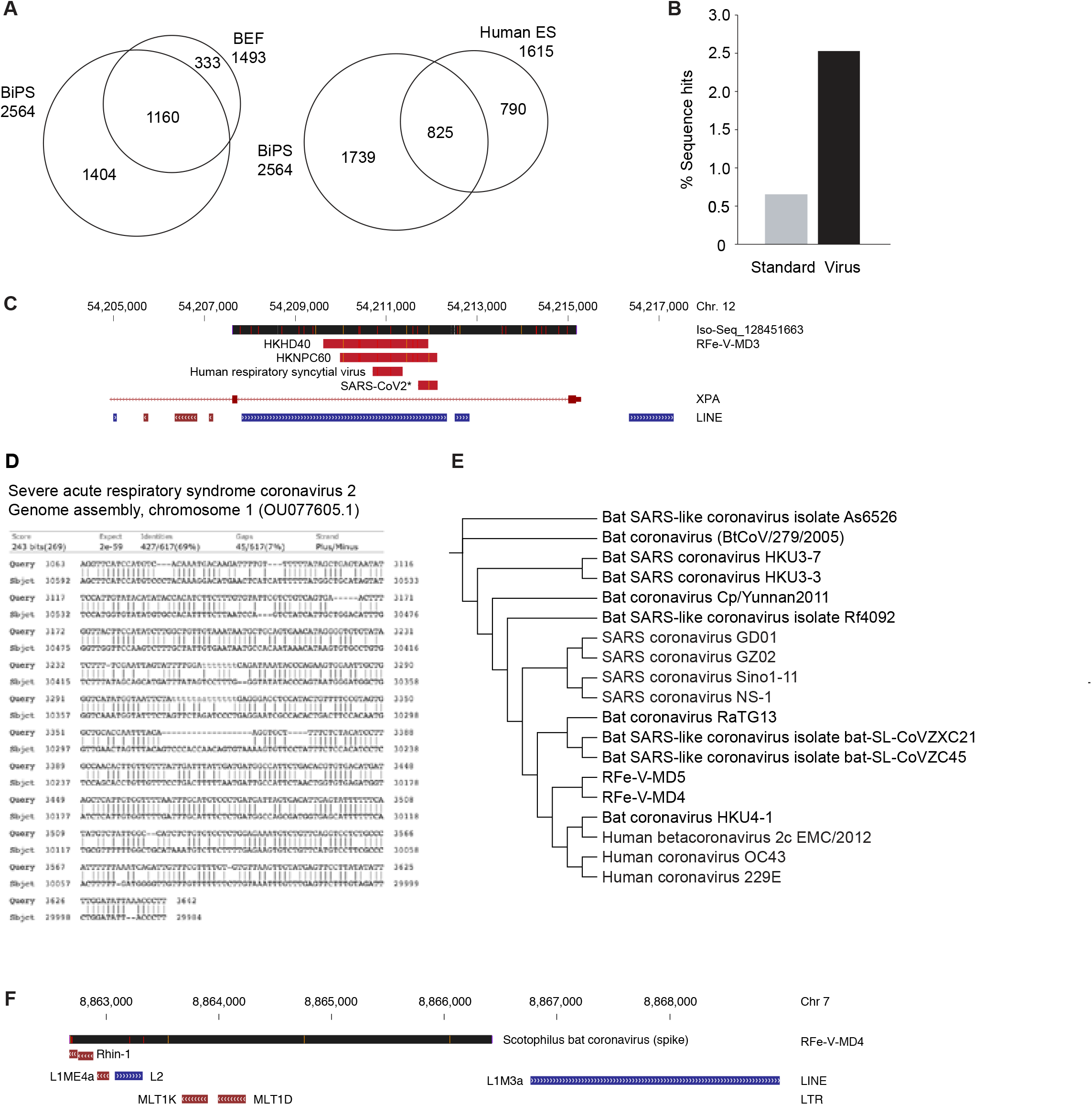
Virome mining in BiPS cells. **(A)** Venn diagram showing the overlap of viral species in human ES cells (GSM4192140 and GSM4192141) and *Rhinolophus ferrumequinum* BiPS cells (top) and BiPS and BEF cells as classified by Kraken2 in RNA-seq reads (bottom). **(B)** Comparison of sequence hits using the Standard or a virus-specific database as reference in Kraken2 analysis with BiPS RNA-seq reads. **(C)** Illustration of a short viral integration with homology to two human herpesvirus 4 isolates (HKD40 and HKNPC60), human respiratory syncytial virus (Kilifi isolate), and a ∼500 bp fragment that was identified at the end of a SARS-CoV2 isolate from an infected patient (NCBI ref. sequence OU077605.1). Shown is the alignment of the 7955 bp-long Iso-seq fragment with the *Rhinolophus ferrumequinum* genome. **(D),** Sequence alignment of the Iso-seq read from BiPS shown in (c) (RFe-V-MD3) with the ∼500 bp SARS-CoV2 fragment described in (C). **(E)** Phylogenetic relationship of the newly identified sequences RFe-V-MD4 and RFe-V-MD5 to a selection of corona viruses. **f,** Genome track of the 6404 bp-long sequence (RFe-V-MD4) with homology to the *Scotophilus* bat coronavirus 512 containing the spike coding region.

**Figure S11.**
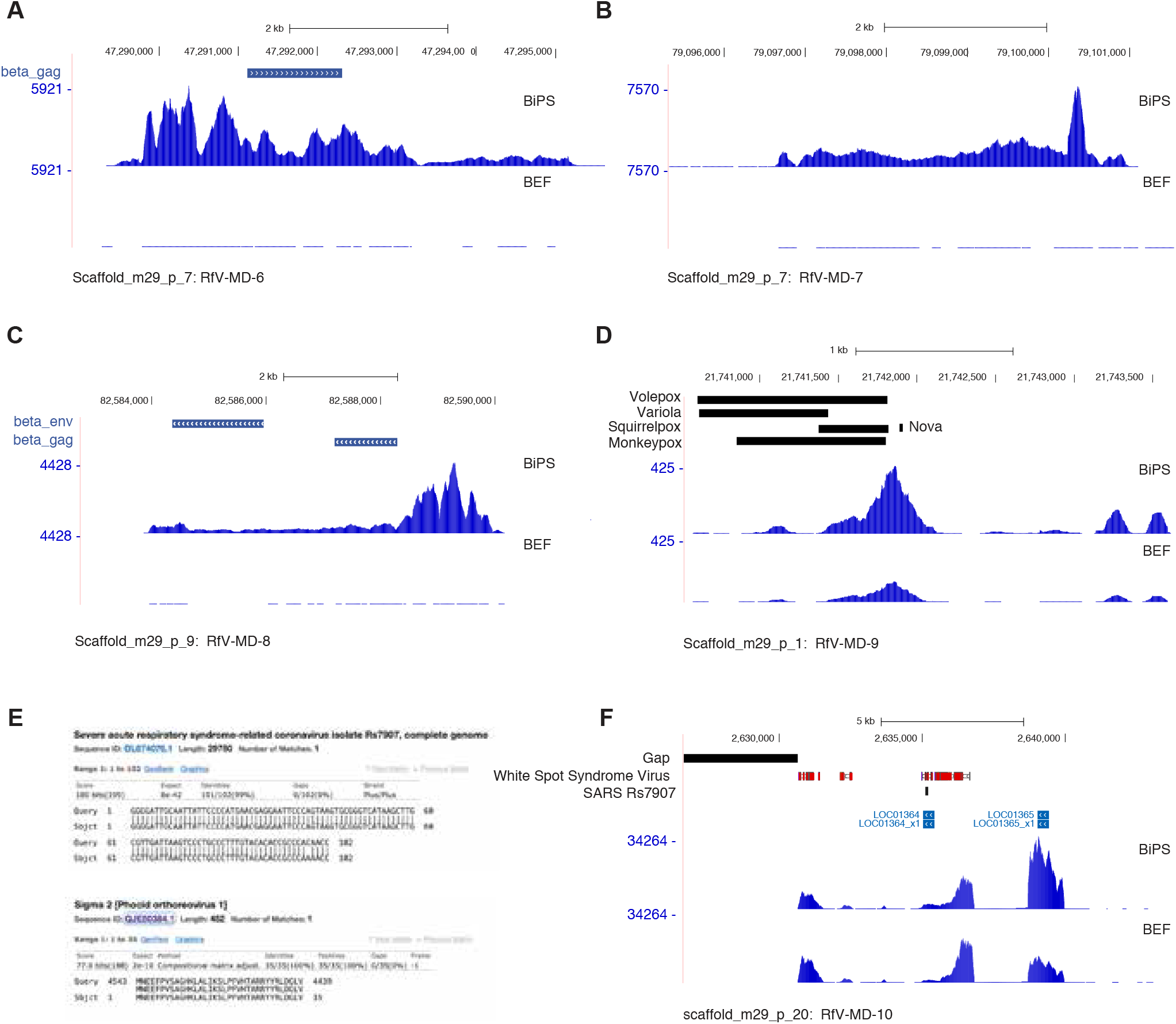
Virome mining in bat iPS and fibroblast cells. Chromosomal regions within indicated *Rhinoophus ferrumequinum* scaffolds detected to contain homologies to the **(A)** Mason-Pfizer monkey virus, the **(B)** Jaagsiekte sheep retrovirus, or the **(C)** Simian endogenous retrovirus. Shown is the surrounding region covered by mapped RNA-seq reads that show further homologies to known viruses on DNA and protein levels as outlined in Table S7. **(D)** Chromosomal region in the vicinity of DNA sequences with homology to several DNA viruses covered by RNA-seq reads that show homologies to cowpox and monkeypox virus proteins after translation using BLASTX. See also Tables S7A and S7G). **(E)** Short homology of expressed sequence reads with *Severe acute respiratory syndrome-related coronavirus isolate Rs7907* on DNA level (*top*) which translates in to an N-terminal fragment of a dsRNA-binding protein (*bottom*). See also Table S7. **(F)** The chromosomal region surrounding the fragment shown in (e) that is covered by mapped sequence reads with further homologies to the White spot syndrome virus. See also Tables S7A and S7H.

**Figure S12.**
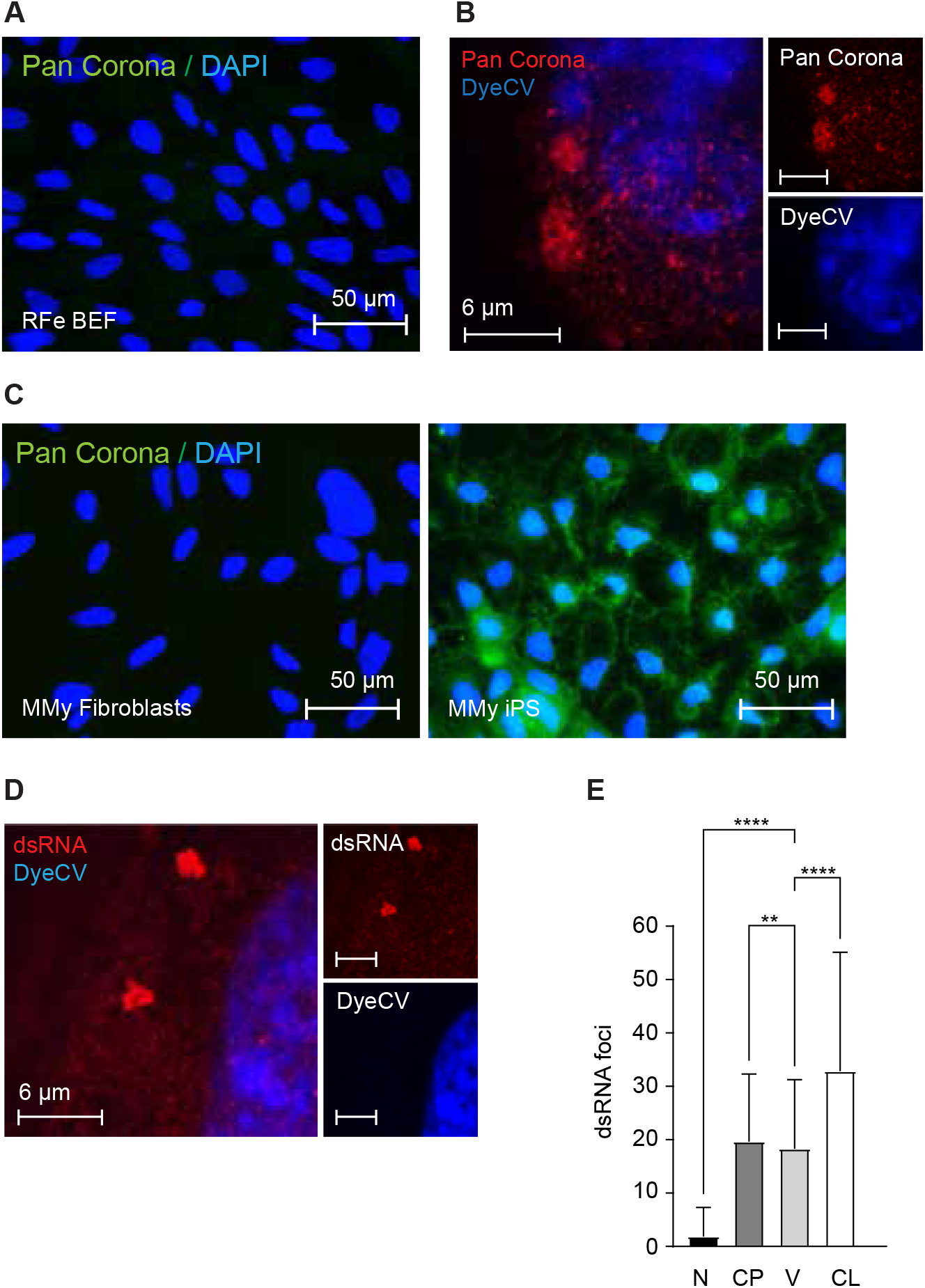
Viral activity in bat pluripotent stem cells. **(A)** Immunostaining with an antibody detecting a Corona antigen in embryonic *Rhinolophus ferrumequinum* embryonic fibroblasts. **(B)** Representative STED microscopy image of bat iPS cells after detecting the Corona antigen as in (A) and DyeCycle Violet (DyeCV) nuclear counter stain and STED microscopy. **(C)** Immunostaining of *Myotis myotis* fibroblasts or induced pluripotent stem cells (iPS) detecting a corona antigen as in (A). **(D)** Representative STED microscopy image of *R. ferrumequinum* iPS cells after immunofluorescende staining of double stranded RNA (dsRNA). **(E)** Quantification of dsRNA foci by ImageStream in *R. ferrumequinum* iPS cells. (dsRNA abundance per cell 32.82 ± 22.31). Further the dsRNA aggregated in subcellular bodies (vesicles 18.28±12.96). **P<0.01,****P<0.0001 by One-way ANOVA with Bonferroni’s multiple comparisons test. Data are presented as mean +/-SD; n=1846 cells. RFe, *Rhinolophus ferrumequinum;* BEF, bat embryonic fibroblasts; CP, cytoplasm; CL, cell; iPS, induced pluripotent stem cells; MMy, *Myotis myotis,* N, nucleus; V, vesicle.

**Figure S13.**
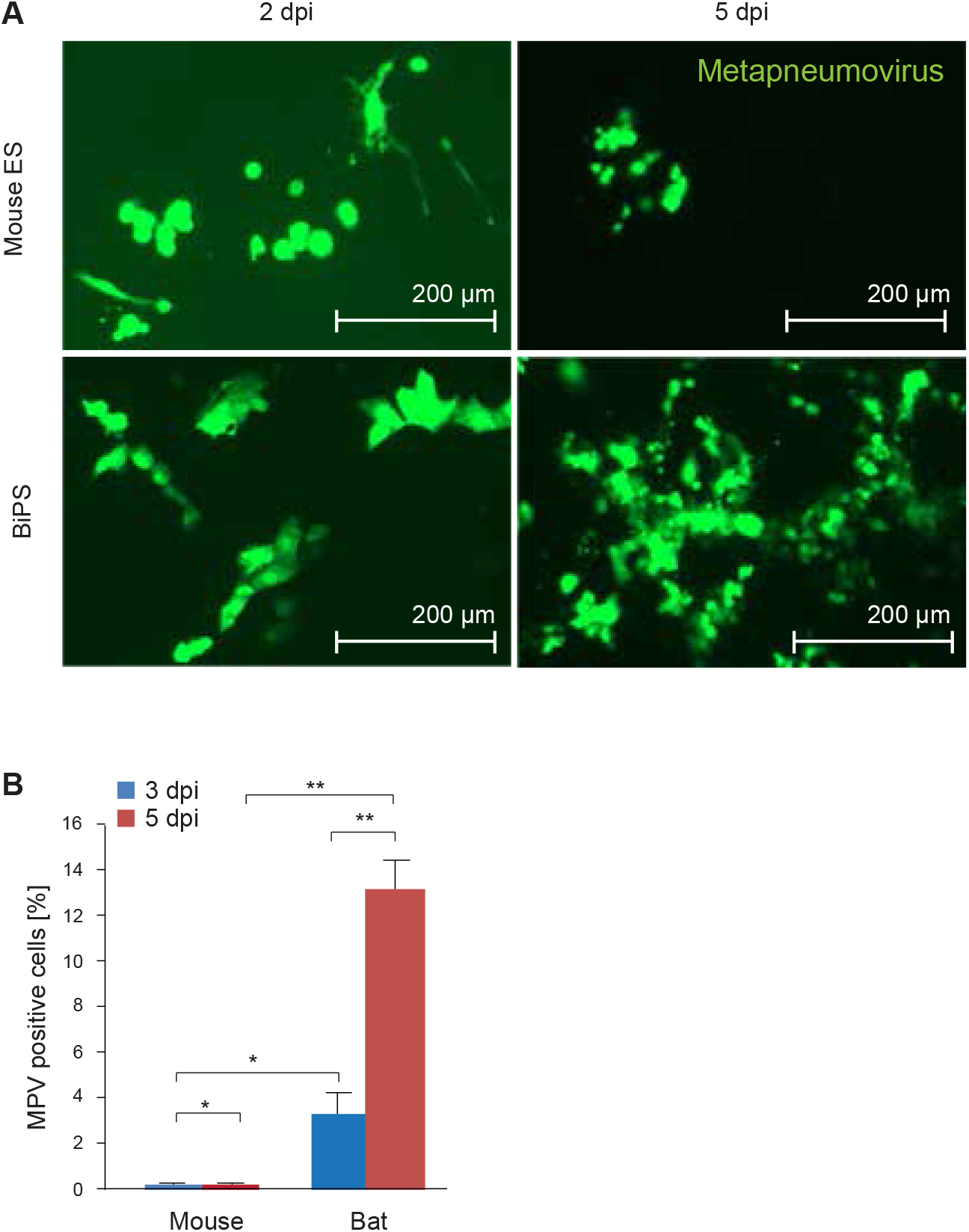
Metapneumovirus infection of mouse and bat pluripotent stem cells. **(A)** Microscopic images of R1 mouse embryonic stem (ES) cells and *Rhinolophus ferrumequinum* induced pluripotent stem cells (BiPS) at 2 and 5 days post-infection (dpi) with GFP-labeled Metapneumovirus (MPV) particles. **(B)** Quantification of experiment shown in (A) at 3 and 5 days post-infection (dpi) by FACS. Data are shown as mean +/-SD; *P<0.05,**P<0.001.

